# Facultatively intra-bacterial localization of a planthopper endosymbiont as an adaptation to its vertical transmission

**DOI:** 10.1101/2023.11.13.566800

**Authors:** Anna Michalik, Diego C. Franco, Teresa Szklarzewicz, Adam Stroiński, Piotr Łukasik

## Abstract

Transovarial transmission is the most reliable way of passing on essential nutrient-providing endosymbionts from mothers to offspring. However, not all endosymbiotic microbes follow the complex path through the female host tissues to oocytes on their own. Here we demonstrate an unusual transmission strategy adapted by one of the endosymbionts of the planthopper *Trypetimorpha occidentalis* (Hemiptera: Tropiduchidae) from Bulgaria. In this species, an *Acetobacteraceae* endosymbiont is transmitted transovarially within deep invaginations of cellular membranes of an ancient endosymbiont *Sulcia* - strikingly resembling recently described plant virus transmission. However, in males, *Acetobacteraceae* colonizes the same bacteriocytes as *Sulcia* but remains unenveloped. Then, the unusual endobacterial localization of *Acetobacteraceae* observed in females appears to be a unique adaptation to maternal transmission. Further, symbiont’s genomic features, including encoding essential amino acid biosynthetic pathways and very similar to a recently described psyllid symbiont, suggest a unique combination of ability to horizontally transmit among species and confer nutritional benefits. The close association with *Acetobacteraceae* symbiont correlates with the so-far- unreported level of genomic erosion of ancient nutritional symbionts of this planthopper. In *Sulcia*, this is reflected in substantial changes in genomic organization, reported for the first time in the symbiont renown for its genomic stability. In *Vidania*, substantial gene loss resulted in one of the smallest genomes known, at 109 kb. Thus, the symbionts of *T. occidentalis* display a combination of unusual adaptations and genomic features that expand our understanding of how insect-microbe symbioses may transmit and evolve.

**Significance Statement:** Reliable transmission across host generations is a major challenge for bacteria that associate with insects, and independently established symbionts have addressed this challenge in different ways. The facultatively endobacterial association of *Acetobacteraceae* symbiont, enveloped by cells of ancient nutritional endosymbiont *Sulcia* in females but not males of the planthopper *Trypetimorpha occidentalis*, appears to be a unique adaptation to maternal transmission. Acetobacteraceae’s genomic features indicate its unusual evolutionary history, and the genomic erosion experienced by ancient nutritional symbionts demonstrates apparent consequences of such close association. Combined, this multi-partite symbiosis expands our understanding of the diversity of strategies that insect symbioses form and some of their evolutionary consequences.

## Introduction

Many insects owe their evolutionary success to symbiosis with microorganisms that influence many aspects of their biology, including nutrition, development, and resistance to various environmental and biotic challenges (1–3). In particular, associations with diverse microorganisms have played a crucial role in the evolution of insect adaptation to nutrition- poor diets, such as phloem or xylem saps and vertebrate blood. Insects that exclusively consume such unbalanced food maintain within their tissues intracellular bacterial or fungal symbionts that supplement the diet with limiting nutrients - essential amino acids and vitamins (4). Some of these symbiotic relationships date back tens or hundreds of millions of years and show a high level of host-microbe integration manifested by metabolic dependency resulting from symbionts’ genome reduction and horizontal gene transfer between partners (5). These ancient symbionts are characterized by extremely reduced genomes, resulting from a massive loss of genes relative to the putative ancestor, likely exceeding 95% in many cases, reaching a stable state where only essential genes remain. Such extreme genome erosion may facilitate the acquisition of new, more versatile symbionts. However, despite the differences in the number or identity of nutritional symbionts reported from diverse sap-feeding insects, we almost always see convergence in the overall amino acid and sometimes vitamin biosynthetic capacity of the symbiotic consortium (6, 7).

The conservation in the function of independently evolved multi-partner symbioses is achieved through complementarity. Generally, when symbionts with overlapping nutritional functions establish a stable infection in the same host, one of the redundant copies of each gene becomes pseudogenized and eliminated. These processes lead to the division of functions required from symbiosis among different symbionts. For example, in different clades of Auchenorrhyncha, *Sulcia* and its independently acquired co-symbionts share responsibilities for the production of 10 essential amino acids. Depending on the Auchenorrhyncha clade, *Sulcia* provides three, seven, or eight amino acids, whereas its partner synthesizes the remainder (8, 9). High level of integration and complementarity was also achieved by endobacterial symbioses of mealybugs from subfamily Pseudococcinae (10, 11). In this case, *Tremblaya* and gammaproteobacteria residing in its cytoplasm have established an interdependent metabolic patchwork, partitioning nutrient biosynthesis. For example, genes involved in the phenylalanine biosynthesis pathway are scattered among the genomes of *Tremblaya*, Gammaproteobacteria, and/or host (11–13). The long-term evolution of such patterns requires reliable symbiont transmission across host generations. Insects and their symbionts have evolved various strategies of symbiont transmission, but the most reliable in the long run is transovarial transmission through the infestation of female germ cells, ensuring the presence of symbionts in each subsequent generation (14–16). This transmission strategy has been adopted many times independently, although the specific mechanisms vary. For example, the ancestral nutritional endosymbionts of all Auchenorrhyncha subfamilies are transmitted transovarially in the same way, namely infecting the posterior end of the ovariole (i.e. structural unit of the ovariole). Some of the more recently acquired endosymbionts follow the same transmission strategy, but others utilize different approaches and infect undifferentiated germ cells or young oocytes (7, 16, 17). It is clear that for any recent arrivals, establishing a reliable transmission means is crucial for maintaining symbiosis. However, depending on the existing functions or pre-adaptations of both the symbionts and the hosts, different strategies may be available.

Here, we present an unusual symbiotic system, that of a Tropiduchidae planthopper *Trypetimorpha occidentalis* from Bulgaria, where an *Acetobacteraceae* symbiont may become enveloped by, and transmitted within, the cells of an ancient symbiont, *Sulcia*. We used a combination of microscopy and genomics to describe the transmission strategy and genomic features of this unusual, facultative *Sulcia* associate, understand its origin and biology, and describe how it may have enabled considerable genome erosion in its co-symbionts.

## Results

### Sex-determined facultatively intra-bacterial localization of *Acetobacteraceae* symbiont

Light microscopy, transmission electron microscopy, and fluorescence *in situ* hybridization have indicated that all examined males and females of *T. occidentalis* are associated with the same set of bacterial endosymbionts: *Sulcia*, *Vidania*, and bacteria from *Acetobacteraceae* family.

Ancient symbionts *Sulcia* and *Vidania* occupy distinct bacteriomes, similar to other planthopper species examined so far (7, 18) (Fig 1). Bacteriomes harboring *Vidania* are syncytial and surrounded by a very thin, monolayered, epithelial bacteriome sheath (Fig. 1A). Their mitochondria-rich cytoplasm is tightly packed with large, multi-lobed *Vidania* cells (Fig. 1A-C). In females, besides lobed *Vidania* localized in the bacteriocytes in the body cavity, we observed the second *Vidania* morphotype that occupies the bacteriocytes in the rectal organ (not shown; see (18)).

**Figure 1.**
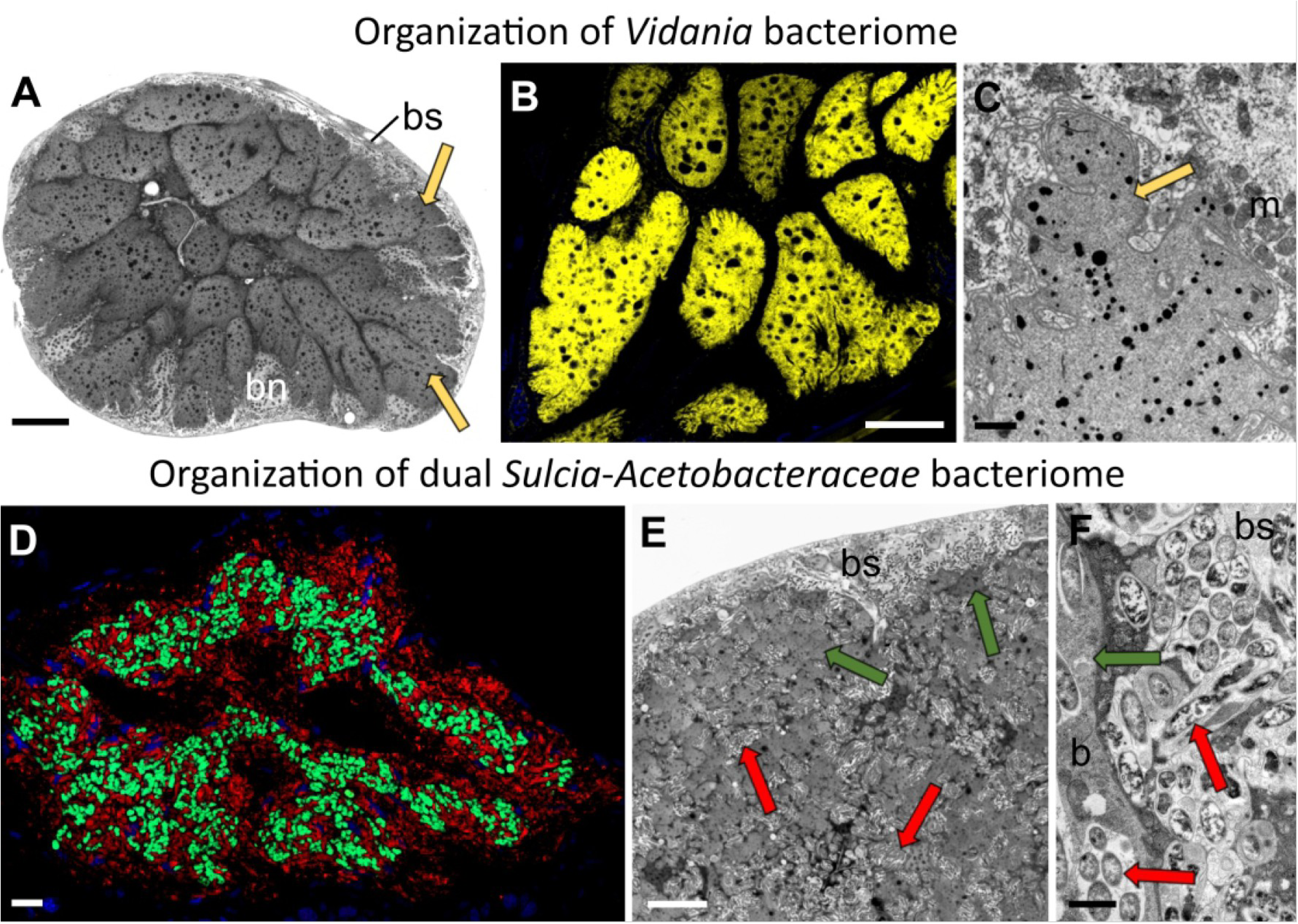
The general overview of the symbiont localization in *T. occidentalis* tissues. **A.** The organization of the *Vidania* bacteriome. LM, scale bar - 10 μm. **B.** Visualisation of *Vidania* cells using *Vidania*-specific probe (yellow), CM, scale bar - 10 μm. **C.** Ultrastructure of *Vidania* cell. TEM, scale bar - 1 μm. **D.** The organization of dual *Acetobacteraceae*-*Sulcia* bacteriome visualized using symbiont-specific probes. Green indicates *Sulcia*, and red indicates *Acetobacteraceae* symbionts. scale bar - 1 μm. **E.** Fragment of the bacteriome harboring *Sulcia* and *Acetobacteraceae* symbionts. LM, scale bar - 10 μm. **F.** *Acetobacteraceae* symbionts in the cytoplasm of bacteriome sheaths. Arrows indicate symbionts: green - *Sulcia*; yellow - *Vidania*; red - *Acetobacteraceae* symbiont; bn - bacteriome nucleus; bs - bacteriome sheath; m – mitochondria

The second ancient planthopper symbiont, *Sulcia*, resides within distinct bacteriomes composed of several bacteriocytes and covered by a thick bacteriome sheath, which creates the invaginations into the bacteriomes (Fig. 1D-F). However, unlike in most other planthoppers, it shares a bacteriome with the *Acetobacteraceae* symbiont. Pleomorphic *Sulcia* cells occur in the bacteriocytes only. In contrast, rod-shaped *Acetobacteraceae* are scattered across the whole bacteriome: they occur in bacteriome sheaths, in the sheath’s invaginations between bacteriomes, and are intermixed with *Sulcia* cells in the cytoplasm of the bacteriocytes (Fig. 1D-F). Their cells may be distributed individually or form clusters, with several cells surrounded by a common external membrane (Fig. 2B, C, F). We did not find *Acetobacteraceae* cells in the gut lumen, where it has been observed in other insects (not shown).

**Figure 2.**
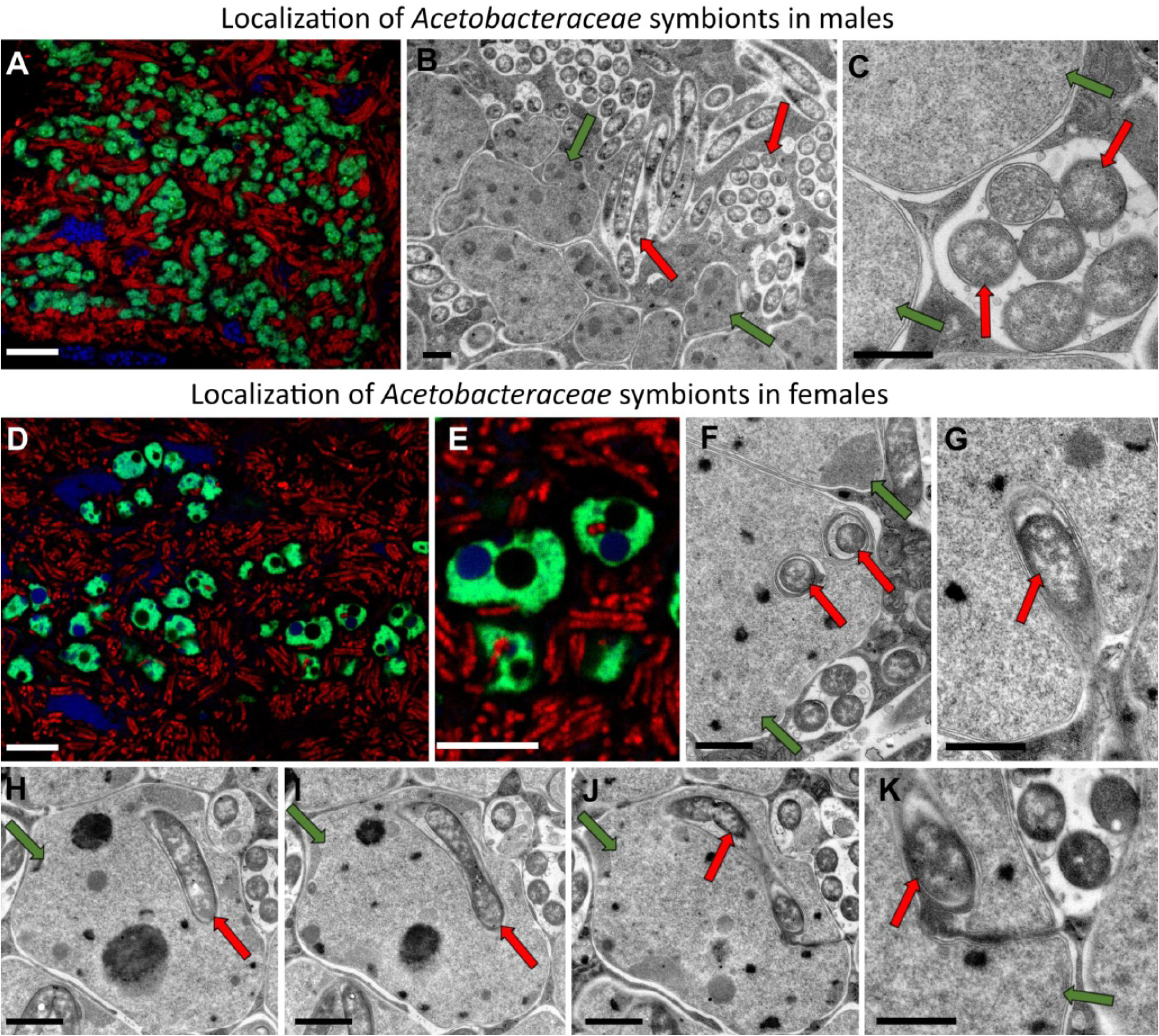
Localization of *Acetobacteraceae* symbionts in male and female tissues. **A.** FISH detection of *Acetobacteraceae* symbionts in the bacteriocytes in males. Green - *Sulcia*, red - *Acetobacteraceae* symbionts. **B, C.** Ultrastructure of *Sulcia* and *Acetobacteraceae* symbionts. TEM, scale bar - 1 μm. **D, E.** Localization of *Acetobacteraceae* symbionts in females. **F-K.** Intra-*Sulcia* localization of *Acetobacteraceae* symbiont within the bacteriome; serial section through the same *Sulcia* cell (H-K) show that the *Acetobacteraceae* remains connected with bacteriocyte cytoplasm through cytoplasmic channels. TEM, scale bar - 1 μm. Arrows indicate symbionts: green - *Sulcia*; red - *Acetobacteraceae* symbiont

While the general organization of the dual *Sulcia*-*Acetobactereaceae* bacteriome is the same in males and females, we found a key difference among sexes in *Acetobacteraceae* localization (Fig. 2A-D). In females only, some of the *Acetobacteraceae* cells within a bacteriocyte are enveloped by the *Sulcia* cells (Fig. 2D-K). They initially appeared to be entirely contained within *Sulcia* cells; however, serial sections indicated that they are always connected to the bacteriocyte cytoplasm by a narrow channel (Fig. 2J, K). In males, we have never observed such envelopment: *Sulcia* and *Acetobacteraceae* were always separate (Fig. 2B, C).

### *Acetobacteraceae* symbionts are transmitted transovarially within *Sulcia* cells

Histological and ultrastructural observations of serial sections showed that all endosymbionts associated with *T. occidentalis* are transovarially transmitted among generations. All symbionts are transmitted simultaneously during the late vitellogenesis stage of oocyte development. The general pattern of transmission of *Sulcia* and *Vidania* is the same as in other planthoppers (7). Both symbionts leave the bacteriocytes and move towards the ovaries. Then, they invade the posterior end of the ovarioles containing fully grown oocytes, migrating to the perivitelline space through the cytoplasm of follicular cells surrounding the posterior pole of the oocyte. Within some of the *Sulcia* cells, at all stages of the migration, we observe *Acetobacteraceae* symbiont cells. We have never observed *Acetobacteraceae* symbionts migrating through the follicular cells on their own; they seem to migrate exclusively when enveloped by *Sulcia* cells (Fig. 3A, B). Toward the end of the migration, all three symbionts gather in the perivitelline space and form a symbiotic ball. Initially, *Acetobacteraceae* symbionts within symbiotic ball are still enveloped by *Sulcia* cells (Fig. 3C, E). However, as oogenesis progresses and egg envelopes thicken, *Acetobacteraceae* cells seem to be separating from *Sulcia* cells (Fig. 3D).

**Figure 3.**
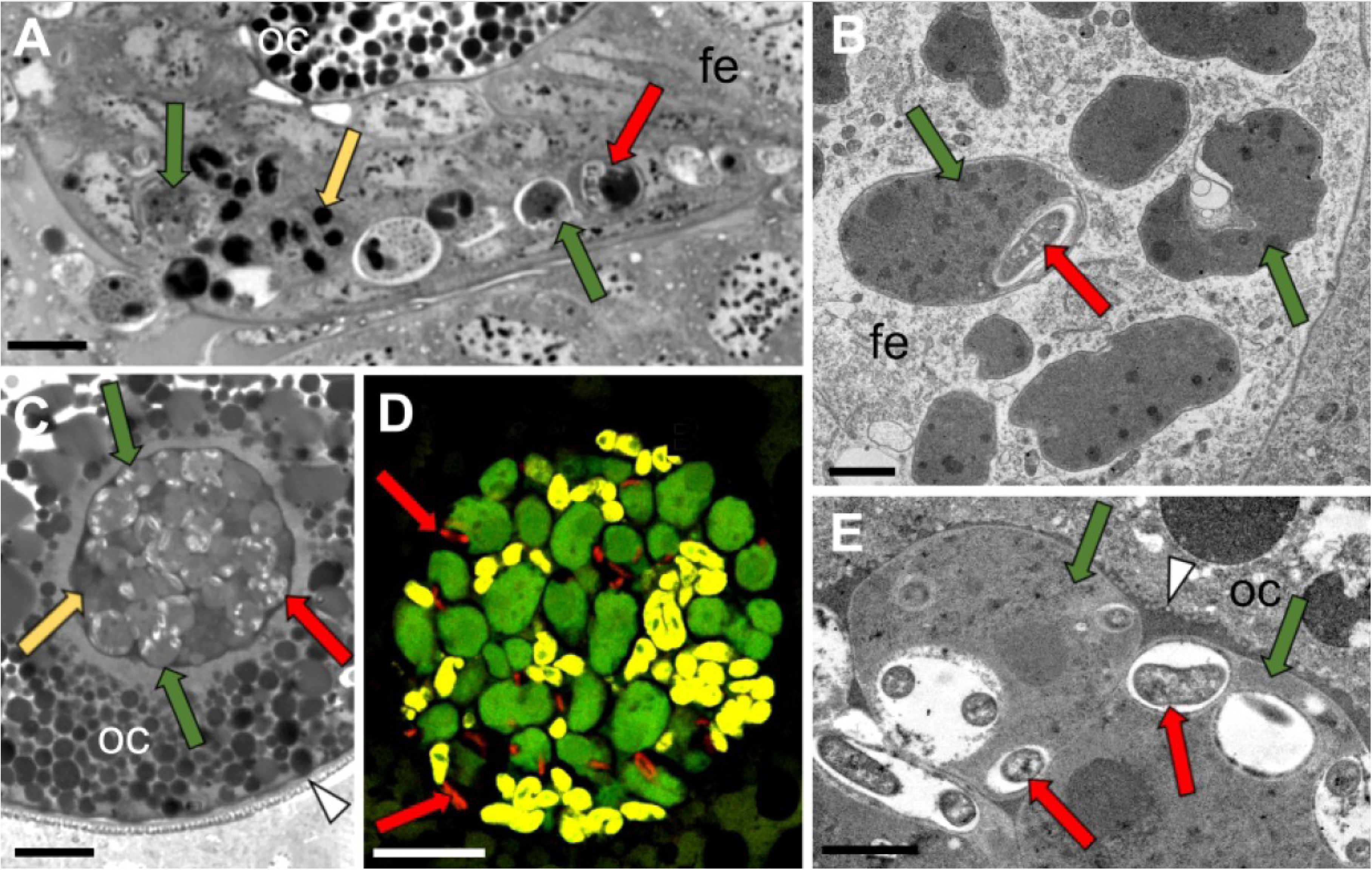
The transovarial transmission of *T. occidentalis* symbionts. A-B. Symbionts migrating through the cytoplasm of follicular cells surrounding the posterior pole of the terminal oocyte. **A.** LM, scale bar - 10 μm. **B.** TEM, scale bar - 1 μm. **C.** A symbiont ball containing bacteria *Sulcia*, *Vidania* and *Acetobacteraceae*, localized in the deep invagination of the oocyte membrane. LM, scale bar - 10 μm. **D.** FISH identification of symbionts in the symbiont ball in mature oocyte. Note that *Acetobacteraceae* cells are at the margins or outside of *Sulcia* cells. CM, scale bar - 10 μm. **E.** Fragment of the symbiont ball in the periviteline space. TEM, scale bar - 1 μm. Arrows indicate symbionts: green - *Sulcia*; yellow - *Vidania*; red - *Acetobacteraceae* symbiont; arrowhead - egg envelopes; fe - follicular epithelium; oc - oocyte.

### Metagenomics-based characterization of *T. occidentalis* microbiota

In order to verify the identity of symbionts associated with *T. occidentalis* and establish their genomic characteristics, roles in symbiosis, and mutual complementation pattern, we sequenced the bacteriome metagenome of a single female (labeled as TRYOCC, from the first letters of genus and species names). In the assembly, contigs representing bacteria differed in their collective size, GC%, coverage, and taxonomic annotation (Fig. 4A). The analysis revealed the presence of three bacterial symbionts: *Vidania* (*Betaproteobacteria*), *Sulcia* (*Bacteroidetes*), and the third one, assigned to *Alphaproteobacteria*. We have fully assembled all of their genomes (Fig. 4B-D).

**Figure 4.**
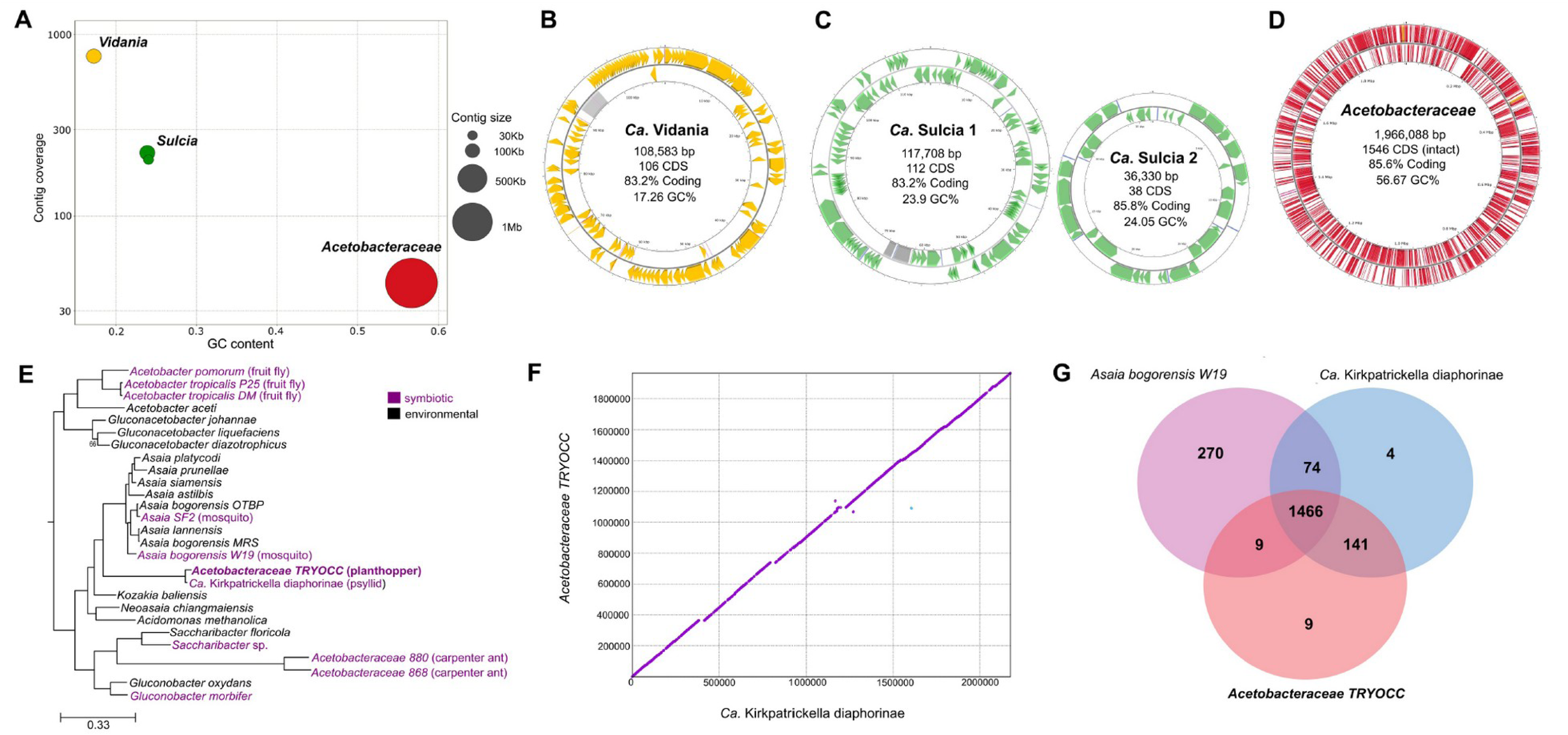
Genomic features of heritable symbionts of *T. occidentalis.* **A.** Metagenome-assembled symbiont genomes plotted in GC contents-coverage space, with contig size represented by circle size **B-D.** Visualizations of *Vidania*, *Sulcia*, and *Acetobacteraceae* genomes with basic genome characteristics. Note that *Sulcia* is represented by two circular chromosomes with similar GC% and coverage and non-overlapping gene sets, which we believe represent a single genome. **E.** Maximum-likelihood of *Acetobacteraceae* phylogeny based on 171 conserved single-copy protein-coding genes, rooted with *Rhodospirillum* sp. as an outgroup. Nodes had bootstrap values of 100, unless otherwise indicated. Animal-associated *Acetobacteraceae* are highlighted using purple font, with host species in the parentheses. Accession numbers of genomes used for phylogenomic analysis are listed in Table S3. **F.** The co-linearity between the genomes of *Ca.* Kirkpatrickella diaphorinae and the *Acetobacteraceae* symbiont of *T. occidentalis*, based on protein space alignment using promer **G.** Venn diagram showing the shared and unique orthologous groups found in *Asaia bogorensis* W19 (mosquito), *Ca.* Kirkpatrickella diaphorinae, and the *Acetobacteraceae* symbiont of *T. occidentalis*.

*Vidania* genome exhibits a size of 108,583 bp, with GC contents of 17.26%, and encodes 106 CDS (Fig. 4B). For *Sulcia*, we obtained two circular contigs with non-overlapping gene sets, of 117,224 bp and 36,330 bp, encoding 112 and 38 CDS, and with GC contents of 23.9% and 24.05%, respectively (Fig. 4C). The alphaproteobacterial symbiont had a circular a genome of 1,966,088 bp and GC% of 56.67, encoding 1546 intact CDS (Fig. 4D).

### *Acetobacteraceae* as heritable endosymbionts of insects

Phylogenomic analyses of the alphaproteobacterial symbiont placed it confidently within the family Acetobacteraceae, revealing its high relatedness to a recently discovered symbiont of a psyllid *Diaphorina citri* from Hawaii - *Candidatus* Kirkpatrickella diaphorinas (hereafter *Kirkpatrickella*) (19). Together, these two strains form a highly supported long-branched clade divergent from other Acetobacteraceae and sister to the genus *Asaia* (Fig 4E).

The genomic comparison between *T. occidentalis’s Acetobacteraceae* symbiont and *Kirkpatrickella* revealed remarkable similarities. They are similar in genome size (1,966,088 vs 2,176,471 bp), nucleotide sequence (average nucleotide identity 92.37%), and genome organization, being perfectly co-linear (Fig. 4F, S1). However, their genomes vary in coding potential and the extent of pseudogenization. TRYOCC symbiont genome includes 1546 intact protein-coding genes, whereas *Kirkpatrickella* contains 1855. Moreover, in the genome of the *Acetobacteraceae* symbiont of *T. occidentalis*, Pseudofinder identified an order of magnitude more pseudogenes than in the symbiont of *D. citri* (831 vs 70), explaining the discrepant number of initial predicted CDS (2295 vs 1909) compared to the genome size (1,966,088 vs 2,176,471 bp) in analyzed symbionts, as pseudogenes may generate the identification of additional non-functional open reading frames.

A functional comparison of *Acetobacteraceae* symbionts revealed a high degree of similarity between the two strains. Both strains had complete pathways related to the biosynthesis of 7 out of 10 essential amino acids (arginine, lysine, phenylalanine, tryptophan, threonine, leucine, and valine), four non-essential amino acids (cysteine, glycine, proline, and serine), as well as two B vitamins (riboflavin and thiamine). Both strains also encode a partial methionine biosynthesis pathway, lacking two genes (*metA*, *metB*) - both absent also in extracellular *Asaia* symbionts of mosquitos and most Auchenorrhyncha symbioses. Interestingly, both *Acetobacteraceae* symbionts retained some genes involved in lipopolysaccharides biosynthesis, pathogenicity (Type 1 secretion system), and drug resistance (Table S1). The presence of these pathways, generally absent in long-term intracellular symbionts, may indicate a relatively early stage of symbiotic association.

The differences in functional gene content between *Acetobacteraceae*-TRYOCC and *Kirkpatrickella* are mainly related in their biosynthetic capacities and include the biosynthesis pathway for two essential amino acids: histidine and isoleucine, and four B vitamins: biotin, cobalamin, folate and pyridoxine. Most of these pathways are complete in the *Kirkpatrickella* genome and incomplete in the Acetobacteraceae symbiont of *T. occidentalis* due to the lack of some genes or their pseudogenization. For example, *Kirkpatrickella* is able to produce histidine and isoleucine, whereas in *Acetobacteraceae*-TRYOCC, three genes in histidine (*hisACE*) and one in isoleucine (*ilvA*) biosynthesis pathways have undergone pseudogenization. Likewise, in the case of B vitamins, incompleteness of their biosynthesis pathways in *Acetobacteraceae*-TRYOCC genome results both from gene loss and pseudogenization. Namely, the symbiont of *T. occidentalis* lost two genes in pyridoxine (*epd* and *pdxB*) and cobalamin (*cobAP*) pathways, whereas two genes involved in biotin biosynthesis (*bioCD*) lost their functionality due to the pseudogenization. Both symbionts have incomplete folate biosynthesis pathway, but they retained different genes: *folE*, *nudB*, *folB*, *folC,* and *folA* in *Acetobacteraceae*-TRYOCC, and *folE*, *folK,* and *folP* in *Kirkpatrella*. We also detected a complete loss of some metabolic pathways in *Acetobacteraceae*-TRYOCC that were present in *Kirkpatrickella* genome, including melatonin biosynthesis, methanogenesis, and proline and adenine degradation (Table S1).

We found more differences in comparisons against other, more distantly related *Acetobacteraceae* (Fig. 4G). For example, the genome of *Asaia bogorensis* from mosquito (Bioproject PRJNA427835) is much larger (genome size 3.9 Mb) and shares 1475 of 1819 orthogroups with TRYOCC symbiont but also encodes 344 orthogroups absent in *T. occidentalis* (Fig. 4G). For instance, the mosquito symbiont possesses genes involved in metabolic pathways, including trehalose biosynthesis, acetate synthesis from acetyl-CoA, and pyrimidine and lysine degradation, that are absent in intracellular *Acetobacteraceae* symbiont of *T. occidentalis* and in *Kirkpatrickella* (Table S1).

### Degenerative changes in ancient nutritional symbiont genomes may be enabled by endobacterial symbiosis

The tiny genomes of *Vidania* and *Sulcia* symbionts of planthoppers have been shown to be very stable in organization and contents over an estimated 200 my of co-diversification with hosts (7, 9, 20). Symbionts of *T. occidentalis* contrast with these patterns: compared to all previously published strains from three planthopper families, both have experienced a series of gene losses and in the case of *Sulcia*, organization changes.

The most striking feature of *Sulcia*-TRYOCC was the substantial change in the genome organization relative to previously characterized genomes. All 82 *Sulcia* genomes published so far in the NCBI database comprised a single circular chromosome and were co- linear relative to each other, except for a single ancestral inversion between Cicadomorpha and Fulgoromorpha and a few additional cases of inversions in different clades (20). In contrast, the genome of *Sulcia*-TRYOCC comprises two distinct circles, both of which have experienced multiple rearrangements relative to previously characterized genomes (Fig. 5A, S2). Genes from different pathways seem to be randomly scattered across these two chromosomes. In contrast, *Vidania-*TRYOCC is co-linear with previously sequenced strains, but the genome comparisons indicate that many genes have been lost. This resulted in the smallest bacterial genome that is not a part of a multi-lineage symbiotic complex (Fig. 5B).

**Figure 5.**
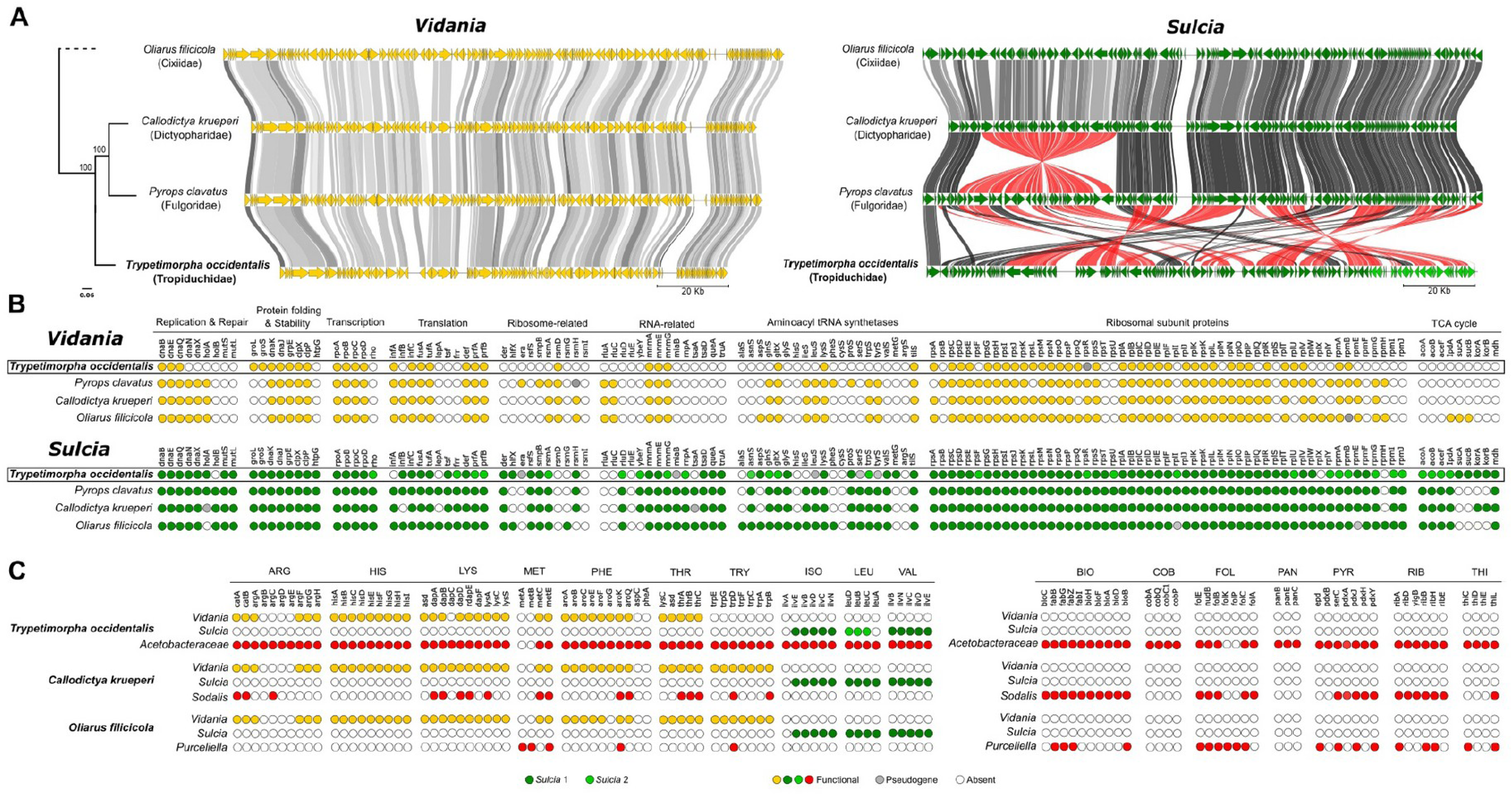
Genomic comparison between symbionts of *T. occidentalis* and representative symbionts from other planthopper families. **A.** Comparative synteny plot of *Vidania* and *Sulcia* genomes. Each circular genome is represented linearly, starting from the *tufA* gene in the case of *Vidania* and *lipB* for *Sulcia*. Arrows indicate genes. Lines connect homologous genes, with red shades indicating inverted genome regions, and line color - the nucleotide sequence similarity. Relationships among host planthopper species, shown to the left of the panel, are based on mitochondrial genomes - redrawn from (18). **B.** Retention of genes in selected functional categories and pathways among *Vidania* and *Sulcia* strains associated with *T. occidentalis* and with representative species from planthopper families Cixiidae, Fulgoridae, and Dictyopharide. Each dot indicates one gene. **C.** Amino acid and B vitamin biosynthesis gene distribution among genomes of different symbionts in *T. occidentalis, C. krueperi*, and *O. filicicola*. In panels B-C, colored dots represent genes present in the genome, grey dots - recognizable pseudogenes, and white - genes that were not detected. In all panels, for *Sulcia* from *T. occidentalis*, genes/regions representing distinct genomic circles are shown using different shades of green.

Compared to *Sulcia* strains representative for planthopper families Fulgoridae and Dictyopharidae (*Sulcia*-PYRCLA and *Sulcia*-CALKRU), *Sulcia*-TRYOCC retained all genes involved in the Krebs cycle but lost some genes from other functional categories. The gene loss is most significant and visible in the aminoacyl-tRNA synthetases group as *Sulcia-* TRYOCC do not possess 11 out of 20 genes (four genes more than other *Sulcia*). Additionally, it lost also single genes involved in DNA repair and replication (*holB*) and translation (infA, lepA) that were present in other *Sulcia* genomes in planthoppers (Fig. 4B). The changes in *Sulcia*-TRYOCC genome did not include genes involved in the biosynthesis of essential amino acids: *Sulcia*-TRYOCC, similarly to *Sulcia* in other planthoppers, provides its host insect with three amino acids, isoleucine, leucine and valine (Fig. 4C).

*Vidania* from TRYOCC has lost more genes relative to *Vidania*-CALKRU and *Vidania*-PYRCLA. The genome reduction involved all functional categories of genes, but particularly notable are the losses in aminoacyl-tRNA synthetases and RNA-related genes. However, the most important from a functional point of view may be the loss of all genes in the tryptophan biosynthesis pathway, reducing the biosynthetic capacity of *Vidania*- TRYOCC. These losses, at least within amino acid biosynthetic pathways, seem to have been enabled by co-symbiosis with *Acetobacteraceae*. All functional genes lost from *Vidania* are present in the *Acetobacteraceae* genome (Fig. 5C).

## Discussion

Decades of research on auchenorrhynchan symbioses have highlighted the remarkable conservation of their ancient symbioses, primarily focused on essential nutritional functions such as amino acid synthesis (4, 21). However, there are several striking exceptions from that stability (22, 23). In the first characterized member of the family Tropiduchidae, several aspects of symbioses stand out among those of other Auchenorrhyncha - or any other known insect nutritional endosymbiotic systems.

To our knowledge, *Acetobacteraceae* have not been reported as endocellular symbionts of insects, but the strain that infects *T. occidentalis* displays the full range of features of an established nutritional endosymbiotic mutualist. Its unique method of transovarial transmission - using other symbiont cells as vessels for transmission across host generations - is, to the best of our knowledge, a phenomenon never before reported from symbiotic bacteria. Its genomic evolutionary features - ongoing pseudogenization of a large share of genes and gradual loss of functions, but without changes in genome organization, contrast with reports of turbulent degenerative processes in other recently acquired symbionts, generally representing Gammaproteobacteria (11, 24). In the sections below, we discussed these unique features, alongside the symbiont’s putative role in the loss of genomic stability in ancient symbionts *Sulcia* and *Vidania*.

### Facultative endobacterial localization as a unique adaptation to transovarial transmission

Transovarial transmission of symbionts between generations ensures the stability of symbiotic interactions. Typically, in Auchenorrhyncha, ancient nutritional endosymbionts transmit across generations in a conserved manner (25, 26). However, newly acquired microorganisms may adopt diverse approaches - likely balancing their own biological features and any pre-adaptations within the host. We demonstrated this in the planthopper family Dictyopharidae, where independently acquired *Sodalis*-allied symbionts have adopted alternative transovarial transmission pathways, colonizing opposite poles of oocytes at different stages of oogenesis (7). Immune system of the insect host, targetting unrecognized “new” microbes within hemolymph, shapes the process of transmission. One strategy to evade an immunological response involves masking surface antigens through co-transmission with “old” symbionts (15). Such transmission strategy in which one symbiont is conveyed inside another has been described in mealybugs (27) and two leafhopper species: *Macrosteles laevis* and *Cicadella viridis* (17, 28). However, in these species, endobacterial localization of gammaproteobacterial symbionts seems permanent, as these symbionts are localized in the cytoplasm of ancient symbionts both in the bacteriocytes and during transmission. At least in mealybugs, long-term endobacterial residence seems to have been an inherent aspect of co-evolution among endosymbionts, which led to their metabolic complementarity (11).

To the best of our knowledge, the tight envelopment of one bacterium by another, as observed in *T. occidentalis*, has not been reported so far from any biological system. *Acetobacteraceae* symbionts remain within *Sulcia* through all stages of transovarial transmission, to the “symbiont ball” stage in the mature oocyte. This intimate association seem to persist at least until the late vitellogenesis stage, when *Acetobacteraceae* cells begin to separate from *Sulcia* cells. We do not currently have data for juvenile stages of *T. occidentalis*; however, it is likely that the symbionts remain separate throughout juvenile development - as observed in adult males - and only re-associate in mature females. The lack of a similar association between *Acetobacteraceae* symbionts and *Sulcia* in males strongly suggests that the facultative endobacterial localization of *Acetobacteraceae* endosymbionts is an adaptation to transovarial transmission. The separation of *Acetobacteraceae* cytoplasm from *Sulcia* cytoplasm by the total of six membranes further suggests their short-term association: in the well-established mealybugs nested symbiosis, *Tremblaya* and its intracytoplasmatic gammaproteobacterial symbiont are separated from each other by two membranes only (12). The six-membrane barrier likely limits metabolic exchange between the two symbionts; however, if, as we suspect, the envelopment only takes place during transmission - a relatively short phase in the insect life cycle - this may not be a limiting factor.

We are not aware of other cases of simultaneous transmission of two symbiotic bacteria mediated by their temporal association. However, there are documented cases of joint transovarial transmission of symbiotic bacteria and viruses (29, 30). In the leafhopper *Nephotettix cincticeps*, rice draw virus (RDV) uses obligate symbiotic bacteria *Sulcia* and *Nasuia* as conveyance vehicles to enter the oocyte and thus pass to the next generation of its insect vector. Virus particles bind to the outer envelope of *Sulcia* and *Nasuia* through direct interaction between outer capsid proteins and bacterial outer membrane proteins (29, 31). The RDV transmission, in deep invaginations of *Sulcia* membrane, striking resembles the transmission of *Acetobacteraceae* symbionts observed in *T. occidentalis*. While the mechanism behind the envelopment of *Acetobacteraceae* by *Sulcia* remains unclear, we can speculate that similar to the RDV case, direct interactions between the outer membrane proteins of the symbionts may mediate the formation of *Sulcia* membrane invaginations. Our understanding of the molecular and cellular mechanisms of obligatory endosymbiont transmission in Auchenorrhyncha remains limited (32). However, detailed ultrastructural analyses have shown that these symbionts pass through the follicular epithelium via the endo-exocytotic pathway, a process with ancient evolutionary origin also employed in the transport of yolk proteins (vitellogenins) from the hemolymph to the oocyte (7). Receptors involved in the transport of vitellogenins into oocytes are known to be used for efficient transmission of facultative endosymbiotic bacteria such as *Wolbachia* or *Spiroplasma* (33, 34). More recently, Mao and co-workers (32) demonstrated that the *Nasuia*-vitellogenin association may facilitate their simultaneous joint entry into host oocytes (32). Hence, it appears that new symbionts adapt to the host’s biology and preferentially utilize the biological mechanisms that are already in place rather than developing entirely new strategies. This adaptive approach can enhance their chances of successful transmission and long-term coexistence with their host.

### Unusual path of an *Acetobacteraceae* strain to heritable nutritional endosymbiosis

Members of the family *Acetobacteraceae* encompass acetous and acidophilic species and are typically found in sugar-rich parts of plants like fruits and flowers, and various products of fermentation. However, recent studies have revealed their versatility as symbionts capable of cross-colonizing phylogenetically diverse insects including ants, honey bees, flies, psyllids, hoppers, cockroaches, and mosquitos (19, 35–38). This broad host spectrum may be attributed to their ability to use multiple transmission routes, including horizontal, venereal, and maternal transmission (35). *Acetobacteraceae* were reported as colonizers of insect digestive tract lumen or salivary glands and, in some cases, of hemolymph and male and female reproductive organs (18, 39). Then, the bacteriocyte-associated *Acetobacteraceae* in *T. occidentalis* may be the first strain convincingly demonstrated to be adapted to insect- endosymbiotic lifestyle, as evidenced by its tissue distribution and genomic characteristics.

Its genome, with a size of 1,966,088 bp, is notably smaller than that of other members of the *Acetobacteraceae* family (except for two carpenter ant symbionts, with similar genome size), and within the range expected for relatively recently acquired nutritional endosymbionts such as *Sodalis*-derived symbionts of other planthoppers (7, 20, 24). However, its comparison with members of the sister clade *Asaia*, and especially with the recently published *Kirkpatrickella* from invasive Hawaiian populations of the psyllid *Diaphorina citri*, suggested genomic evolutionary patterns substantially departing from those observed in better-known gammaproteobacterial symbionts. The first difference is the stability of genome organization, with *Acetobacteraceae* symbiont of *T. occidentalis* fully co-linear with *Kirkpatrickella* and similar to *Asaia* strains. The ongoing genomic degradation process observed in *T. occidentalis Acetobacteraceae* symbiont, evidenced by a large number of pseudogenes that are still functional in *Kirkpatrickella*, seems to be gradual and does not change genome organization. This genomic stability contrasts with the genome evolution pattern observed in more widespread gammaproteobacterial symbionts from *Sodalis* and *Arsenophonus* clades, which tend to undergo rapid genome degradation and rampant rearrangements once they establish within new hosts (24, 40).

*Sodalis*-allied symbionts are thought to be all derived from opportunistic ancestors similar and related to *Sodalis praecaptivus* - free-living bacteria found both in plant and animal tissues including human wound, with a genome size of approximately 5.5Mb (24, 41, 42). In contrast, the phylogenetic proximity and genomic similarity of *T. occidentalis* symbiont and *Kirkpatrickella*, found in phylogenetically distant insects, is indicative of very different biology. Notably, the Hawaiian lineage of *D. citri* must have been colonized relatively recently (as *Kirkpatrickella* is absent in other populations of this globally invasive species), and its symbiont retains the ability to be transmitted through microinjection among individuals (19). While the precise tissue localization of *Kirkpatrickella* in *D. citri* is unknown, it bears characteristics of facultative endosymbionts such as *Wolbachia* - which has been moving within and across species for tens of millions of years (43, 44). However, some *Wolbachia* lineages have established within certain host lineages as obligate nutrient-providing mutualists incapable of shifting hosts again, and this led to further gene loss and genome reduction (45). This is also what may have happened to *T. occidentalis* symbiont, whose gene set is largely a subset of that of *Kirkpatrickella,* with evidence of extensive pseudogenization and ongoing loss of several functions.

However, the key difference between known facultative endosymbionts and the *Kirkpatickella* clade is in functional characteristics. The latter display an impressive nutrient biosynthetic range, whereas facultative endosymbionts such as *Wolbachia* are known primarily for reproductive manipulation and defensive properties, with only some strains known to contribute vitamins (45, 46). Hence, it is tempting to consider the *Kirkpatrickella* clade as a representative of an unusual functional category of “facultative nutritional endosymbionts”, retaining the ability to switch hosts and thus transmit across species. Future work on broader collections of insects may shed further light on the validity of this classification. Nevertheless, it seems that *T. occidentalis* symbiont has evolved into a mutually obligate associate of its planthopper host, with the extent of pseudogenization making it unlikely to retain the capacity to switch hosts again.

### *Acetobacteraceae* infection coincides with departure from genomic stability in ancient nutritional endosymbionts *Sulcia* and *Vidania*

Recently established symbionts usually undergo rapid and turbulent genomic reduction, which slows down as the share of the genome responsible for essential processes increases (24, 47). Ancient nutritional endosymbionts like *Buchnera*, *Sulcia*, and *Vidania* exemplify the remarkable stability of genome organization and contents over tens of millions of years of co-diversification with their insect hosts (20, 48). Genomic rearrangements and loss of functional genes are relatively rare in ancient endosymbionts, with a notable exception of cases of co-infections with additional nutrient-providing symbionts that often lead to complementarity among symbionts in their nutritional and possibly other functions. In *T. occidentalis*, the genomes of both ancient nutritional endosymbionts departed to a large extent from the conserved ancestral state, represented by all other known genomes of planthopper-associated *Sulcia* and *Vidania* strains (7, 9, 20, 49). *Sulcia* symbiont from *T. occidentalis* has experienced multiple rearrangements compared to the ancestral state, comparable to the total number identified from across ∼80 genomes spanning ∼300 my of evolution that have been published so far (20). Even more unusual is the fragmentation of the genome into two distinct genomic circles, or chromosomes.Both genomic rearrangements and genome fragmentation into chromosomes have been reported before, from hemipteran symbionts (23, 50) and organellar genomes (51, 52). The biological significance of these genomic changes is unclear, but at evolutionary timescales, the departure from long-term stability may indicate faster degeneration, potentially speeding up the descent into what Bennett and Moran (47) described as “an evolutionary rabbit hole”.

*Vidania* from *T. occidentalis* retains the ancestral genome organization but it has lost multiple genes and functions relative to strains characterized to date (7, 9, 20). As a result of these changes, *Vidania*-TRYOCC may have the tiniest stand-alone bacterial genome described so far - at 109 kb, smaller than *Nasuia* from leafhoppers (>112 kb) (53), other *Vidania* strains published to date (>122 kb) or any *Sulcia* strains (>142 kb) (7, 9, 20). Tiny genomes of obligatory symbionts of Sternorrhyncha, including *Carsonella* symbiont of psyllids (>160 kb), and *Tremblaya* from mealybugs (>138 kb) are also less reduced (11, 54). An exception are the genomes of interdependent lineages of an alphaproteobacterium *Hodgkinia* that comprise unique multi-lineage symbiotic complexes in some cicadas. Lineages share the ancestral set of ca. 150 genes / 150 kb of the nucleotide sequence, and the tiniest of them may encode fewer than 20 genes on an <80 kb genome, but they require other lineages to ensure basic cellular function (23, 55).

It is tempting to propose that these substantial and unusual genomic changes in both symbionts of *T. occidentalis* have occurred relatively recently, as a result of infection by *Acetobacteraceae*, and its particularly close association with *Sulcia.* The close association and the potential metabolic interactions between these symbionts may have triggered genomic changes and the complementary loss of certain functions in *Sulcia* and *Vidania*. This is exemplified by the absence of the tryptophan operon in the *Vidania* genome, which is apparently complemented by the *Acetobacteraceae* symbiont. This kind of complementarity, where different symbionts present in the same host share responsibilities for essential metabolic functions including tryptophan biosynthesis, has been documented in other multi- partite systems, such as *Buchnera-Serratia* and *Tremblaya-*Gammaproteobacteria complexes (11, 56). Unfortunately, the scarcity of currently available genomic references prevent a comprehensive description of the dynamics of genomic degeneration. Over an estimated >100 million years of evolution separating *T. occidentalis* from the closest planthopper species with characterized symbioses, many changes and adaptations may have occurred, including the possibility of serial replacements of the symbionts associated with *Sulcia* and *Vidania*. Further research and a more comprehensive database of symbiotic associations in planthoppers are required to understand the evolutionary events that have shaped their nutritional endosymbioses.

### Conclusions: Speeding up the discovery of bacterial strategies

The comprehensive characterization of symbioses of the first member of the planthopper family Tropiduchidae revealed unexpected results regarding new lifestyles adopted by members of a well-known bacterial family, new types of interactions and associations among bacteria, added to the knowledge of functional categories of insect symbionts, and the evolution and stability of symbiont genomes.

During nearly three decades since the publication of the first bacterial genome (57), it is clear that we have not progressed very far in uncovering what Paul Buchner (1965) famously described as “the veritable fairyland of insect symbiosis”. Given the microbial symbionts’ importance in insect biology, function, and environmental adaptation, at scales ranging from individual life history traits to population and community processes (58, 59), it is critical to comprehensively characterize these patterns as global biodiversity declines (60, 61).

## Material and Methods

### Study material

The specimens of the planthopper *Trypetimorpha occidentalis* Huang & Bourgoin, 1993 originate from a single population sampled in Harsovo, Bulgaria, in July 2018. After collection, insects were identified based on morphological features and preserved whole in ethanol or partially dissected and fixed in a 2.5% glutaraldehyde solution. After fixation, they were stored until use at 4 °C.

### Microscopic analyses

#### Histological and ultrastructural analyses

The dissected abdomens of the adult males and females were fixed in 2.5% glutaraldehyde in 0.1 M phosphate buffer (pH 7.2) at 4°C. The fixed material was then rinsed three times in the same buffer with the addition of sucrose (5.8 g/100 ml) and postfixed in 1% osmium tetroxide for 2 hours at room temperature. After postfixation, samples were dehydrated in a graded series of ethanol (30-100%) and acetone, embedded in epoxy resin Epon 812 (Merck, Darmstadt, Germany), and cut into sections using Reichert-Jung ultracut E microtome. Semithin sections (1 µm thick) were stained in 1% methylene blue in 1% borax and analyzed and subsequently photographed under a Nikon Eclipse 80i light microscope (LM). Ultrathin sections (90 nm thick) were contrasted with uranyl acetate and lead citrate and examined and photographed in Jeol JEM 2100 electron transmission microscope (TEM) at 80 kV.

#### Fluorescence in situ hybridization

Fluorescence *in situ* hybridization was performed using fluorochrome-labelled oligonucleotide probes targeting 16S rRNA of symbionts associated with *T. occidentalis* (Table S2). Insects preserved in ethanol were rehydrated and then postfixed in 4% paraformaldehyde for two hours at room temperature. Next, the material was dehydrated again by incubation in increased concentrations of ethanol (30-100%) and acetone, embedded in Technovit 8100 resin (Kulzer, Wehrheim, Germany), and cut into semithin sections (1 um thick). The sections were then incubated overnight at room temperature in a hybridization buffer containing the specific sets of probes with a final concentration of 100 nM. After hybridization, the slides were washed in PBS three times, dried, covered with ProLong Gold Antifade Reagent (Life Technologies), and examined using a confocal laser scanning microscope Zeiss Axio Observer LSM 710 (CM).

### Metagenomic library preparation and sequencing

We sequenced bacteriome metagenomic library for *T. occidentalis* (TRYOCC). DNA from dissected bacteriomes of individual female, extracted using the Sherlock AX kit (A&A Biotechnology, Gdynia, Poland), was fragmented using a Covaris E220 sonicator and used for metagenomic library preparation using the NEBNext Ultra II DNA Library Prep kit for Illumina (New England BioLabs), with the target insert length of 350 bp. The library pool, including three target species and other samples, was sequenced on an Illumina HiSeq X SBS lane by NGXBio (San Francisco, CA, USA).

### Metagenome characterization and symbiont genome annotation

Raw sequencing reads were quality filtered and had adapters removed using Trim Galore v0.6.4 (settings: –length 80 -q 30; https://github.com/FelixKrueger/TrimGalore). High-quality filtered reads were checked using FastQC v0.11.9 (https://github.com/s-andrews/FastQC). Contigs were assembled using Megahit v1.2.9, with a maximum k-mer size = 255 and min contig size = 1,000 (62) Symbiont contigs were identified using NanoTax.py (https://github.com/diecasfranco/Nanotax). This python script performs blast searches against a customized nucleotide database and a protein database containing sequences from previously assembled genomes of hosts, symbionts, and their free-living relatives. Other information, including the GC content, coverage, and length of each contig, is compiled into output with the assigned taxonomy.

The *Acetobacteraceae* symbiont contigs were extracted and mapped against the raw reads. A new *de novo* assembly using extracted reads was performed using SPAdes v.3.14.1, with a kmer list (-k 55,77,99,127) and –isolate option (63). The assembly generated two contigs and were manually curated and merged into a circular genome. Lower-coverage regions and gaps were checked using Tablet v.1.21.02.08 (64). The genome of *Vidania* assembled into a single circularly mapping contig. The genome of *Sulcia* assembled into two circularly mapping contigs, with no indication of alternative arrangements such as large numbers of irregularly mapping reads.

The circular genome of *Acetobacteraceae* symbiont was annotated using bakta v1.8.1 (65) with default parameters. Protein fasta files obtained from bakta pipeline were submitted to GhostKOALA (66) for metabolic pathways’ reconstruction. Pseudogenes were identified using pseudofinder (67). The genomic contigs of *Sulcia* and *Vidania* were annotated with a custom Python script modified from (23). The script predicts all the Open Reading Frames (ORFs) and their amino acid sequences from each genome. These ORFs were searched recursively using HMMER v3.3.1 (68) against custom databases containing manually curated sets of protein-coding, rRNA, and noncoding RNA (ncRNA) genes from previously characterized *Sulcia* or *Vidania* lineages. rRNA and ncRNA genes were searched with nhmmer (HMMER V3.3.1) (69), and tRNAs were identified with tRNAscan-SE v2.0.7 (70). Based on the relative length compared to the reference genes, protein-coding genes were classified as functional (>85%), putative pseudogenes (>60%), or pseudogenes (<60%).

The taxon-annotated GC-coverage plots for symbiont contigs were drawn using R v. 4.0.2 (R Development Core Team) with the ggplot2 package. Genomes were visualized using Proksee (71). Comparative synteny plots were obtained using pyGenomeViz package(https://github.com/moshi4/pyGenomeViz).

Phylogenomic analysis from the *Acetobacteraceae* clade was performed by extracting the single copy genes detected among whole genomes by BUSCO v 5.4.3 (72), using the *Rhodospirillales* lineage model. Individual alignments for each BUSCO genes were performed using MUSCLE v3.8.1551 (73). The alignments were concatenated using seqkit tool (74). IQ-tree was used to infer the phylogenetic tree based on the best substitution model according to ModelFinder (LG+F+R5). Bootstrapping was conducted using “SH-aLRT” bootstrap (BS) methods with 1,000 replicates. All other setting options were set as default.

The orthologus gene clusters from Acetobacteraceae symbionts of *T. occidentalis*, *D. citri* and *Asaia bogorensis W19* were obtained using OrthoVenn3 (75), with the option orthoMCL and default parameters.

## Acknowledgments

We thank John McCutcheon for permission to use the facilities at the University of Montana and Ada Jankowska for laboratory assistance.

## Author Contributions

AM, DCF, PŁ designed research; AS provided the insect samples; AM, DCF, TS, PŁ performed research and analyzed data; AM, DCF, PŁ wrote the manuscript. All authors reviewed the manuscript and approved the final version.

## Data availability statement

Sequence data have been deposited in GenBank under Bioproject PRJNA1031153. Additional results, including Pseudofinder results and bioinformatic pipelines, are available in the GitHub repository (https://github.com/diecasfranco/TRYOCC_metagenomics).

