## Supplementary information for "Facultatively intra-bacterial localization of a planthopper endosymbiont as an adaptation to its vertical transmission"

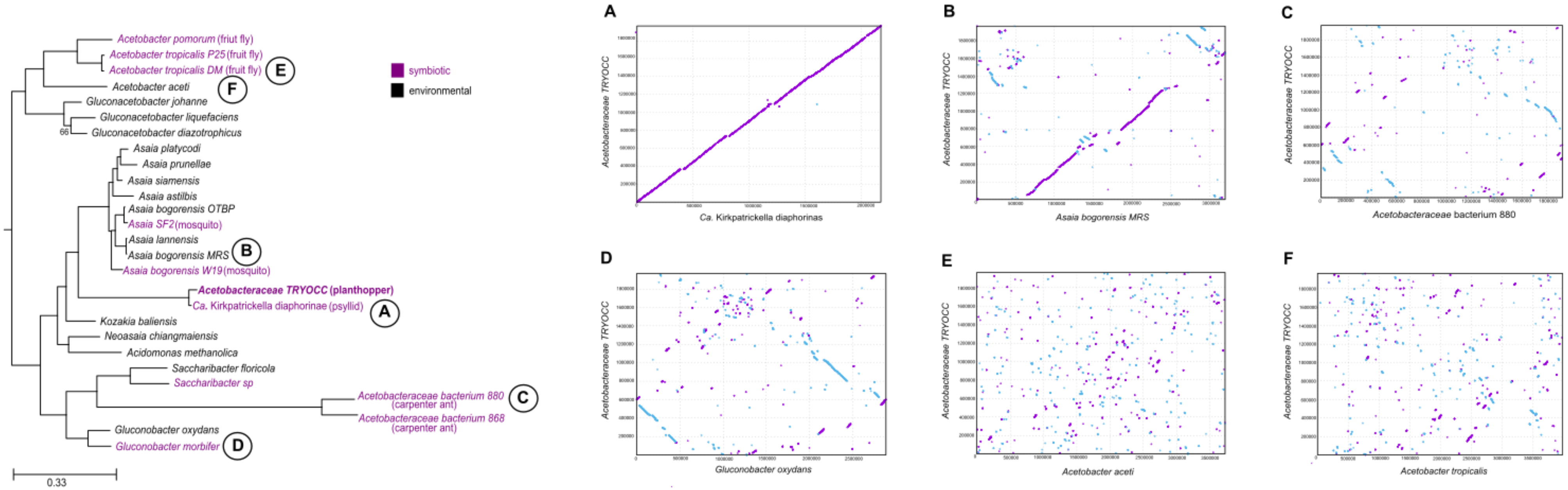

**Figure S1.** PROMER alignments of *Acetobacteraceae TRYOCC* genome against the genomes of various representatives of Acetobacteraceae family: **A.** *Ca.* Kirkpatrickella diaphorinae B. *Asaia bogorensis MRS* **C.** *Acetobacteraceae bacterium 880* **D.** *Gluconobacter oxydans* **E.** *Acetobacter aceti* **F.** *Acetobacter tropicalis*. The bacterial phylogeny and panel A overlap with Figure 4.

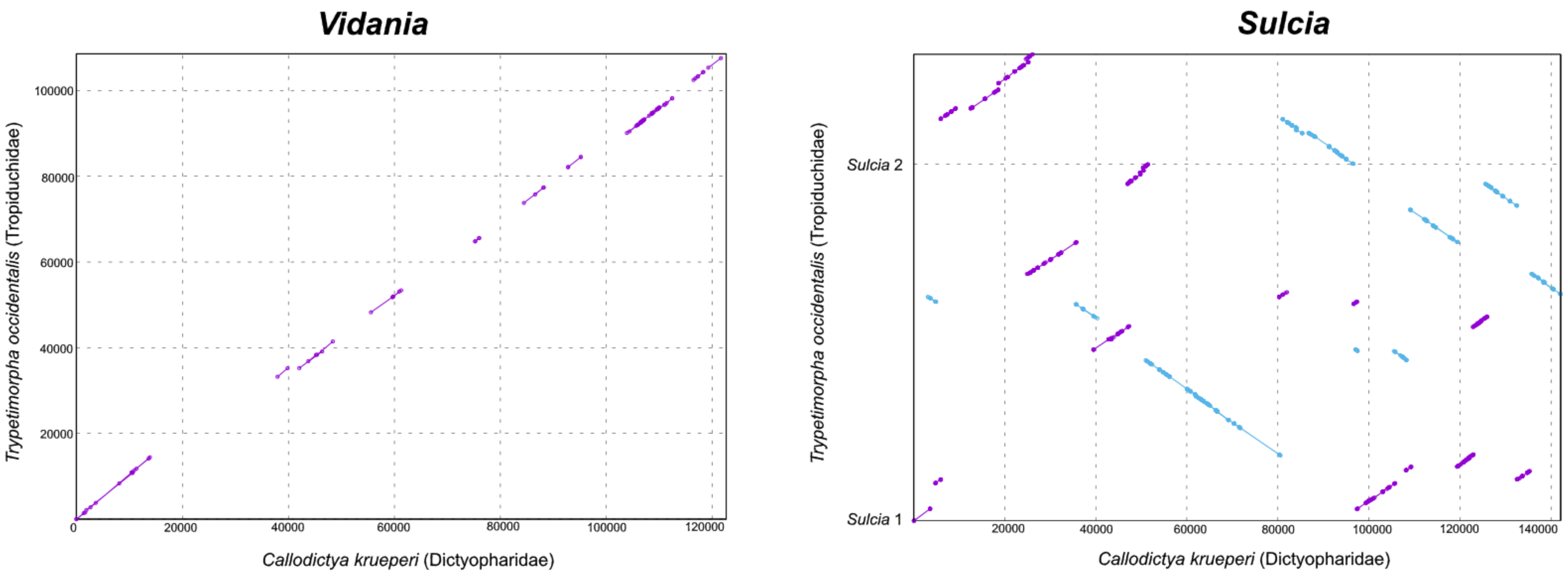

**Figure S2.** PROMER alignments of *Sulcia* and *Vidania* genomes from *T. occidentalis* (Tropiduchidae) against genomes from *Callodictya krueperi* (Dictyopharidae).

| *Asaia bogorensis_W19* | *Ca.* **Kirkpatrella diaphorinas** | *Acetobacteraceae TRYOCC* |
|---|---|---|
| **Carbohydrate metabolism** | **Carbohydrate metabolism** | **Carbohydrate metabolism** |
| Central carbohydrate metabolism | Central carbohydrate metabolism | Central carbohydrate metabolism |
| M00001 Glycolysis (Embden-Meyerhof pathway), glucose => pyruvate (10) (1 block missing 8/9) | M00001 Glycolysis (Embden-Meyerhof pathway), glucose => pyruvate (9) (2 blocks missing 7/9) | M00001 Glycolysis (Embden-Meyerhof pathway), glucose => pyruvate (9) (2 blocks missing 7/9) |
| M00002 Glycolysis, core module involving three-carbon compounds (6) (complete 5/5) | M00002 Glycolysis, core module involving three-carbon compounds (5) (1 block missing 4/5) | M00002 Glycolysis, core module involving three-carbon compounds (5) (1 block missing 4/5) |
| M00003 Gluconeogenesis, oxaloacetate => fructose-6P (8) (1 block missing 6/7) | M00003 Gluconeogenesis, oxaloacetate => fructose-6P (7) (2 blocks missing 5/7) | M00003 Gluconeogenesis, oxaloacetate => fructose-6P (7) (2 blocks missing 5/7) |
| M00307 Pyruvate oxidation, pyruvate => acetyl-CoA (5) (complete 1/1) | M00307 Pyruvate oxidation, pyruvate => acetyl-CoA (4) (complete 1/1) | M00307 Pyruvate oxidation, pyruvate => acetyl-CoA (4) (complete 1/1) |
| M00009 Citrate cycle (TCA cycle, Krebs cycle) (10) (2 blocks missing 6/8) | M00009 Citrate cycle (TCA cycle, Krebs cycle) (11) (2 blocks missing 6/8) | M00009 Citrate cycle (TCA cycle, Krebs cycle) (11) (2 blocks missing 6/8) |
| M00010 Citrate cycle, first carbon oxidation, oxaloacetate => 2-oxoglutarate (3) (complete 3/3) | M00010 Citrate cycle, first carbon oxidation, oxaloacetate => 2-oxoglutarate (3) (complete 3/3) | M00010 Citrate cycle, first carbon oxidation, oxaloacetate => 2-oxoglutarate (3) (complete 3/3) |
| M00011 Citrate cycle, second carbon oxidation, 2-oxoglutarate => oxaloacetate (7) (2 blocks missing 3/5) | M00011 Citrate cycle, second carbon oxidation, 2-oxoglutarate => oxaloacetate (8) (2 blocks missing 3/5) | M00011 Citrate cycle, second carbon oxidation, 2-oxoglutarate => oxaloacetate (8) (2 blocks missing 3/5) |
| M00004 Pentose phosphate pathway (Pentose phosphate cycle) (9) (complete 6/6) | M00004 Pentose phosphate pathway (Pentose phosphate cycle) (7) (complete 6/6) | M00004 Pentose phosphate pathway (Pentose phosphate cycle) (7) (complete 6/6) |
| M00006 Pentose phosphate pathway, oxidative phase, glucose 6P => ribulose 5P (4) (complete 2/2) | M00006 Pentose phosphate pathway, oxidative phase, glucose 6P => ribulose 5P (3) (complete 2/2) | M00006 Pentose phosphate pathway, oxidative phase, glucose 6P => ribulose 5P (3) (complete 2/2) |
| M00007 Pentose phosphate pathway, non-oxidative phase, fructose 6P => ribose 5P (5) (complete 4/4) | M00007 Pentose phosphate pathway, non-oxidative phase, fructose 6P => ribose 5P (4) (complete 4/4) | M00007 Pentose phosphate pathway, non-oxidative phase, fructose 6P => ribose 5P (4) (complete 4/4) |
| M00580 Pentose phosphate pathway, archaea, fructose 6P => ribose 5P (2) (1 block missing 1/2) | M00580 Pentose phosphate pathway, archaea, fructose 6P => ribose 5P (2) (1 block missing 1/2) | M00580 Pentose phosphate pathway, archaea, fructose 6P => ribose 5P (2) (1 block missing 1/2) |
| M00005 PRPP biosynthesis, ribose 5P => PRPP (1) (complete 1/1) | M00005 PRPP biosynthesis, ribose 5P => PRPP (1) (complete 1/1) | M00005 PRPP biosynthesis, ribose 5P => PRPP (1) (complete 1/1) |
| M00008 Entner-Doudoroff pathway, glucose-6P => glyceraldehyde-3P + pyruvate (5) (complete 4/4) | M00008 Entner-Doudoroff pathway, glucose-6P => glyceraldehyde-3P + pyruvate (2) (2 blocks missing 2/4) | M00008 Entner-Doudoroff pathway, glucose-6P => glyceraldehyde-3P + pyruvate (2) (2 blocks missing 2/4) |
| M00308 Semi-phosphorylative Entner-Doudoroff pathway, gluconate => glycerate-3P (3) (2 blocks missing 2/4) | M00308 Semi-phosphorylative Entner-Doudoroff pathway, gluconate => glycerate-3P (2) (incomplete 1/4) | M00308 Semi-phosphorylative Entner-Doudoroff pathway, gluconate => glycerate-3P (2) (incomplete 1/4) |
| Other carbohydrate metabolism | Other carbohydrate metabolism | Other carbohydrate metabolism |
| M00014 Glucuronate pathway (uronate pathway) (4) (incomplete 3/7) | M00014 Glucuronate pathway (uronate pathway) (2) (incomplete 2/7) | M00014 Glucuronate pathway (uronate pathway) (2) (incomplete 2/7) |
| M00631 D-Galacturonate degradation (bacteria), D-galacturonate => pyruvate + D-glyceraldehyde 3P (1) (incomplete 1/5) | **ABSENT** | **ABSENT** |
| M00061 D-Glucuronate degradation, D-glucuronate => pyruvate + D-glyceraldehyde 3P (1) (incomplete 1/5) | M00061 D-Glucuronate degradation, D-glucuronate => pyruvate + D-glyceraldehyde 3P (1) (incomplete 1/5) | M00061 D-Glucuronate degradation, D-glucuronate => pyruvate + D-glyceraldehyde 3P (1) (incomplete 1/5) |
| M00632 Galactose degradation, Leloir pathway, galactose => alpha-D-glucose-1P (2) (2 blocks missing 2/4) | M00632 Galactose degradation, Leloir pathway, galactose => alpha-D-glucose-1P (2) (2 blocks missing 2/4) | M00632 Galactose degradation, Leloir pathway, galactose => alpha-D-glucose-1P (1) (incomplete 1/4) |
| M00552 D-galactonate degradation, De Ley-Doudoroff pathway, D-galactonate => glycerate-3P (2) (incomplete 2/5) | M00552 D-galactonate degradation, De Ley-Doudoroff pathway, D-galactonate => glycerate-3P (2) (incomplete 2/5) | M00552 D-galactonate degradation, De Ley-Doudoroff pathway, D-galactonate => glycerate-3P (2) (incomplete 2/5) |
| M00129 Ascorbate biosynthesis, animals, glucose-1P => ascorbate (3) (incomplete 3/7) | M00129 Ascorbate biosynthesis, animals, glucose-1P => ascorbate (3) (incomplete 3/7) | M00129 Ascorbate biosynthesis, animals, glucose-1P => ascorbate (2) (incomplete 2/7) |
| M00854 Glycogen biosynthesis, glucose-1P => glycogen/starch (3) (1 block missing 1/2) | **ABSENT** | **ABSENT** |
| M00855 Glycogen degradation, glycogen => glucose-6P (1) (2 blocks missing 1/3) | M00855 Glycogen degradation, glycogen => glucose-6P (1) (2 blocks missing 1/3) | M00855 Glycogen degradation, glycogen => glucose-6P (1) (2 blocks missing 1/3) |

| | | |
|---|---|---|
| M00565 Trehalose biosynthesis, D-glucose 1P => trehalose (5) (1 block missing 5/6) | ABSENT | ABSENT |
| M00549 Nucleotide sugar biosynthesis, glucose => UDP-glucose (3) (complete 3/3) | M00549 Nucleotide sugar biosynthesis, glucose => UDP-glucose (3) (complete 3/3) | M00549 Nucleotide sugar biosynthesis, glucose => UDP-glucose (3) (complete 3/3) |
| M00892 UDP-N-acetyl-D-glucosamine biosynthesis, eukaryotes, glucose => UDP-GlcNAc (2) (incomplete 2/6) | M00892 UDP-N-acetyl-D-glucosamine biosynthesis, eukaryotes, glucose => UDP-GlcNAc (2) (incomplete 2/6) | M00892 UDP-N-acetyl-D-glucosamine biosynthesis, eukaryotes, glucose => UDP-GlcNAc (2) (incomplete 2/6) |
| M00909 UDP-N-acetyl-D-glucosamine biosynthesis, prokaryotes, glucose => UDP-GlcNAc (5) (complete 5/5) | M00909 UDP-N-acetyl-D-glucosamine biosynthesis, prokaryotes, glucose => UDP-GlcNAc (5) (complete 5/5) | M00909 UDP-N-acetyl-D-glucosamine biosynthesis, prokaryotes, glucose => UDP-GlcNAc (5) (complete 5/5) |
| M00012 Glyoxylate cycle (2) (incomplete 2/5) | M00012 Glyoxylate cycle (2) (incomplete 2/5) | M00012 Glyoxylate cycle (2) (incomplete 2/5) |
| M00740 Methylaspartate cycle (2) (incomplete 2/11) | M00740 Methylaspartate cycle (2) (incomplete 2/11) | M00740 Methylaspartate cycle (2) (incomplete 2/11) |
| M00532 Photorespiration (6) (incomplete 3/10) | M00532 Photorespiration (7) (incomplete 4/10) | M00532 Photorespiration (5) (incomplete 2/10) |
| M00013 Malonate semialdehyde pathway, propanoyl-CoA => acetyl-CoA (1) (incomplete 1/5) | ABSENT | ABSENT |
| M00131 Inositol phosphate metabolism, Ins(1,3,4,5)P4 => Ins(1,3,4)P3 => myo-inositol (1) (incomplete 1/4) | M00131 Inositol phosphate metabolism, Ins(1,3,4,5)P4 => Ins(1,3,4)P3 => myo-inositol (1) (incomplete 1/4) | M00131 Inositol phosphate metabolism, Ins(1,3,4,5)P4 => Ins(1,3,4)P3 => myo-inositol (1) (incomplete 1/4) |
| **Energy metabolism** | **Energy metabolism** | **Energy metabolism** |
| Carbon fixation | Carbon fixation | Carbon fixation |
| M00165 Reductive pentose phosphate cycle (Calvin cycle) (8) (2 blocks missing 9/11) | M00165 Reductive pentose phosphate cycle (Calvin cycle) (7) (incomplete 8/11) | M00165 Reductive pentose phosphate cycle (Calvin cycle) (7) (incomplete 8/11) |
| M00168 CAM (Crassulacean acid metabolism), dark (1) (1 block missing 1/2) | M00168 CAM (Crassulacean acid metabolism), dark (1) (1 block missing 1/2) | M00168 CAM (Crassulacean acid metabolism), dark (1) (1 block missing 1/2) |
| M00172 C4-dicarboxylic acid cycle, NADP - malic enzyme type (1) (incomplete 1/4) | M00172 C4-dicarboxylic acid cycle, NADP - malic enzyme type (1) (incomplete 1/4) | M00172 C4-dicarboxylic acid cycle, NADP - malic enzyme type (1) (incomplete 1/4) |
| M00171 C4-dicarboxylic acid cycle, NAD - malic enzyme type (1) (incomplete 1/7) | M00171 C4-dicarboxylic acid cycle, NAD - malic enzyme type (1) (incomplete 1/7) | M00171 C4-dicarboxylic acid cycle, NAD - malic enzyme type (1) (incomplete 1/7) |
| M00170 C4-dicarboxylic acid cycle, phosphoenolpyruvate carboxykinase type (1) (incomplete 1/4) | M00170 C4-dicarboxylic acid cycle, phosphoenolpyruvate carboxykinase type (1) (incomplete 1/4) | M00170 C4-dicarboxylic acid cycle, phosphoenolpyruvate carboxykinase type (1) (incomplete 1/4) |
| M00173 Reductive citrate cycle (Arnon-Buchanan cycle) (7) (incomplete 3/10) | M00173 Reductive citrate cycle (Arnon-Buchanan cycle) (7) (incomplete 3/10) | M00173 Reductive citrate cycle (Arnon-Buchanan cycle) (7) (incomplete 3/10) |
| M00376 3-Hydroxypropionate bi-cycle (7) (incomplete 2/13) | M00376 3-Hydroxypropionate bi-cycle (8) (incomplete 3/13) | M00376 3-Hydroxypropionate bi-cycle (8) (incomplete 3/13) |
| M00374 Dicarboxylate-hydroxybutyrate cycle (3) (incomplete 1/13) | M00374 Dicarboxylate-hydroxybutyrate cycle (4) (incomplete 2/13) | M00374 Dicarboxylate-hydroxybutyrate cycle (4) (incomplete 2/13) |
| M00377 Reductive acetyl-CoA pathway (Wood-Ljungdahl pathway) (1) (incomplete 1/7) | M00377 Reductive acetyl-CoA pathway (Wood-Ljungdahl pathway) (1) (incomplete 1/7) | M00377 Reductive acetyl-CoA pathway (Wood-Ljungdahl pathway) (1) (incomplete 1/7) |
| M00579 Phosphate acetyltransferase-acetate kinase pathway, acetyl-CoA => acetate (2) (complete 2/2) | ABSENT | ABSENT |
| Methane metabolism | Methane metabolism | Methane metabolism |
| M00357 Methanogenesis, acetate => methane (2) (incomplete 1/5) | M00357 Methanogenesis, acetate => methane (1) (incomplete 1/5) | ABSENT |
| M00346 Formaldehyde assimilation, serine pathway (3) (incomplete 3/9) | M00346 Formaldehyde assimilation, serine pathway (3) (incomplete 3/9) | M00346 Formaldehyde assimilation, serine pathway (3) (incomplete 3/9) |
| M00345 Formaldehyde assimilation, ribulose monophosphate pathway (2) (2 blocks missing 1/3) | M00345 Formaldehyde assimilation, ribulose monophosphate pathway (2) (2 blocks missing 1/3) | M00345 Formaldehyde assimilation, ribulose monophosphate pathway (2) (2 blocks missing 1/3) |
| M00344 Formaldehyde assimilation, xylulose monophosphate pathway (2) (2 blocks missing 2/4) | M00344 Formaldehyde assimilation, xylulose monophosphate pathway (1) (incomplete 1/4) | M00344 Formaldehyde assimilation, xylulose monophosphate pathway (1) (incomplete 1/4) |
| Sulfur metabolism | Sulfur metabolism | ABSENT |
| M00176 Assimilatory sulfate reduction, sulfate => H2S (4) (1 block missing 1/2) | M00176 Assimilatory sulfate reduction, sulfate => H2S (4) (1 block missing 1/2) | |

| | | |
|---|---|---|
| ATP synthesis | ATP synthesis | ATP synthesis |
| M00144 NADH:quinone oxidoreductase, prokaryotes (13) (complete 1/1) | M00144 NADH:quinone oxidoreductase, prokaryotes (13) (complete 1/1) | M00144 NADH:quinone oxidoreductase, prokaryotes (13) (complete 1/1) |
| | M00149 Succinate dehydrogenase, prokaryotes (4) (complete 1/1) | M00149 Succinate dehydrogenase, prokaryotes (4) (complete 1/1) |
| M00417 Cytochrome o ubiquinol oxidase (4) (complete 1/1) | M00417 Cytochrome o ubiquinol oxidase (4) (complete 1/1) | M00417 Cytochrome o ubiquinol oxidase (4) (complete 1/1) |
| **Lipid metabolism** | **Lipid metabolism** | **Lipid metabolism** |
| Fatty acid metabolism | Fatty acid metabolism | Fatty acid metabolism |
| M00082 Fatty acid biosynthesis, initiation (6) (complete 2/2) | M00082 Fatty acid biosynthesis, initiation (6) (complete 2/2) | M00082 Fatty acid biosynthesis, initiation (6) (complete 2/2) |
| M00083 Fatty acid biosynthesis, elongation (5) (complete 1/1) | M00083 Fatty acid biosynthesis, elongation (5) (complete 1/1) | M00083 Fatty acid biosynthesis, elongation (5) (complete 1/1) |
| M00873 Fatty acid biosynthesis in mitochondria, animals (2) (incomplete 1/6) | M00873 Fatty acid biosynthesis in mitochondria, animals (2) (incomplete 1/6) | M00873 Fatty acid biosynthesis in mitochondria, animals (2) (incomplete 1/6) |
| M00874 Fatty acid biosynthesis in mitochondria, fungi (3) (incomplete 2/6) | M00874 Fatty acid biosynthesis in mitochondria, fungi (3) (incomplete 2/6) | M00874 Fatty acid biosynthesis in mitochondria, fungi (3) (incomplete 2/6) |
| Lipid metabolism | Lipid metabolism | Lipid metabolism |
| M00089 Triacylglycerol biosynthesis (1) (incomplete 1/4) | M00089 Triacylglycerol biosynthesis (1) (incomplete 1/4) | M00089 Triacylglycerol biosynthesis (1) (incomplete 1/4) |
| M00091 Phosphatidylcholine (PC) biosynthesis, PE => PC (1) (complete 1/1) | M00091 Phosphatidylcholine (PC) biosynthesis, PE => PC (1) (complete 1/1) | M00091 Phosphatidylcholine (PC) biosynthesis, PE => PC (1) (complete 1/1) |
| M00093 Phosphatidylethanolamine (PE) biosynthesis, PA => PS => PE (3) (complete 3/3) | M00093 Phosphatidylethanolamine (PE) biosynthesis, PA => PS => PE (3) (complete 3/3) | M00093 Phosphatidylethanolamine (PE) biosynthesis, PA => PS => PE (3) (complete 3/3) |
| M00066 Lactosylceramide biosynthesis (1) (1 block missing 1/2) | M00066 Lactosylceramide biosynthesis (1) (1 block missing 1/2) | M00066 Lactosylceramide biosynthesis (1) (1 block missing 1/2) |
| **Nucleotide metabolism** | **Nucleotide metabolism** | **Nucleotide metabolism** |
| Purine metabolism | Purine metabolism | Purine metabolism |
| M00048 De novo purine biosynthesis, PRPP + glutamine => IMP (13) (complete 8/8) | M00048 De novo purine biosynthesis, PRPP + glutamine => IMP (12) (complete 8/8) | M00048 De novo purine biosynthesis, PRPP + glutamine => IMP (12) (complete 8/8) |
| M00049 Adenine ribonucleotide biosynthesis, IMP => ADP,ATP (4) (complete 4/4) | M00049 Adenine ribonucleotide biosynthesis, IMP => ADP,ATP (3) (1 block missing 3/4) complete - missing gene manually found in the genome | M00049 Adenine ribonucleotide biosynthesis, IMP => ADP,ATP (4) (complete 4/4) |
| M00050 Guanine ribonucleotide biosynthesis, IMP => GDP,GTP (4) (complete 4/4) | M00050 Guanine ribonucleotide biosynthesis, IMP => GDP,GTP (3) (1 block missing 3/4) complete - missing gene manually found in the genome | M00050 Guanine ribonucleotide biosynthesis, IMP => GDP,GTP (4) (complete 4/4) |
| M00053 Deoxyribonucleotide biosynthesis, ADP/GDP/CDP/UDP => dATP/dGTP/dCTP/dUTP (3) (complete 2/2) | M00053 Deoxyribonucleotide biosynthesis, ADP/GDP/CDP/UDP => dATP/dGTP/dCTP/dUTP (2) (1 block missing 1/2) complete - missing gene manually found in the genome | M00053 Deoxyribonucleotide biosynthesis, ADP/GDP/CDP/UDP => dATP/dGTP/dCTP/dUTP (3) (complete 2/2) |
| M00958 Adenine ribonucleotide degradation, AMP => Urate (5) (complete 3/3) | M00958 Adenine ribonucleotide degradation, AMP => Urate (2) (1 block missing 2/3) | **ABSENT** |
| M00959 Guanine ribonucleotide degradation, GMP => Urate (5) (complete 4/4) | M00959 Guanine ribonucleotide degradation, GMP => Urate (2) (2 blocks missing 2/4) | M00959 Guanine ribonucleotide degradation, GMP => Urate (1) (incomplete 1/4) |
| M00546 Purine degradation, xanthine => urea (6) (1 block missing 4/5) | M00546 Purine degradation, xanthine => urea (1) (incomplete 1/5) | M00546 Purine degradation, xanthine => urea (1) (incomplete 1/5) |
| Pyrimidine metabolism | Pyrimidine metabolism | Pyrimidine metabolism |
| M00051 De novo pyrimidine biosynthesis, glutamine (+ PRPP) => UMP (7) (1 block missing 2/3) | M00051 De novo pyrimidine biosynthesis, glutamine (+ PRPP) => UMP (7) (1 block missing 2/3) | M00051 De novo pyrimidine biosynthesis, glutamine (+ PRPP) => UMP (7) (1 block missing 2/3) |
| M00052 Pyrimidine ribonucleotide biosynthesis, UMP => UDP/UTP,CDP/CTP (3) (complete 3/3) | M00052 Pyrimidine ribonucleotide biosynthesis, UMP => UDP/UTP,CDP/CTP (2) (1 block missing 2/3) complete - missing gene manually found in the genome | M00052 Pyrimidine ribonucleotide biosynthesis, UMP => UDP/UTP,CDP/CTP (3) (complete 3/3) |
| M00938 Pyrimidine deoxyribonucleotide biosynthesis, UDP => dTTP (6) (complete 5/5) | M00938 Pyrimidine deoxyribonucleotide biosynthesis, UDP => dTTP (5) (1 block missing 4/5) complete - missing gene manually found in the genome | M00938 Pyrimidine deoxyribonucleotide biosynthesis, UDP => dTTP (6) (complete 5/5) |

| | | |
|---|---|---|
| M00046 Pyrimidine degradation, uracil => beta-alanine, thymine => 3-aminoisobutanoate (4) (complete 3/3) | ABSENT | ABSENT |
| M00939 Pyrimidine degradation, uracil => 3-hydroxypropanoate (3) (incomplete 1/5) | M00939 Pyrimidine degradation, uracil => 3-hydroxypropanoate (2) (incomplete 1/5) | M00939 Pyrimidine degradation, uracil => 3-hydroxypropanoate (2) (incomplete 1/5) |
| **Amino acid metabolism** | **Amino acid metabolism** | **Amino acid metabolism** |
| Serine and threonine metabolism | Serine and threonine metabolism | Serine and threonine metabolism |
| M00020 Serine biosynthesis, glycerate-3P => serine (3) (complete 3/3) | M00020 Serine biosynthesis, glycerate-3P => serine (3) (complete 3/3) | M00020 Serine biosynthesis, glycerate-3P => serine (3) (complete 3/3) |
| M00018 Threonine biosynthesis, aspartate => homoserine => threonine (5) (complete 5/5) | M00018 Threonine biosynthesis, aspartate => homoserine => threonine (5) (complete 5/5) | M00018 Threonine biosynthesis, aspartate => homoserine => threonine (5) (complete 5/5) |
| M00621 Glycine cleavage system (3) (complete 3/3) | M00621 Glycine cleavage system (3) (complete 3/3) | M00621 Glycine cleavage system (3) (complete 3/3) |
| M00033 Ectoine biosynthesis, aspartate => ectoine (2) (incomplete 2/5) | M00033 Ectoine biosynthesis, aspartate => ectoine (2) (incomplete 2/5) | M00033 Ectoine biosynthesis, aspartate => ectoine (2) (incomplete 2/5) |
| Cysteine and methionine metabolism | Cysteine and methionine metabolism | Cysteine and methionine metabolism |
| M00021 Cysteine biosynthesis, serine => cysteine (2) (complete 2/2) | M00021 Cysteine biosynthesis, serine => cysteine (2) (complete 2/2) | M00021 Cysteine biosynthesis, serine => cysteine (2) (complete 2/2) |
| M00609 Cysteine biosynthesis, methionine => cysteine (1) (incomplete 1/6) | M00609 Cysteine biosynthesis, methionine => cysteine (1) (incomplete 1/6) | M00609 Cysteine biosynthesis, methionine => cysteine (1) (incomplete 1/6) |
| M00017 Methionine biosynthesis, aspartate => homoserine => methionine (7) (1 block missing 6/7) | M00017 Methionine biosynthesis, aspartate => homoserine => methionine (7) (1 block missing 6/7) | M00017 Methionine biosynthesis, aspartate => homoserine => methionine (5) (2 blocks missing 5/7) |
| M00034 Methionine salvage pathway (10) (1 block missing 7/8) | M00034 Methionine salvage pathway (5) (incomplete 4/8) | M00034 Methionine salvage pathway (4) (incomplete 4/8) |
| M00035 Methionine degradation (3) (1 block missing 3/4) | M00035 Methionine degradation (2) (2 blocks missing 2/4) | M00035 Methionine degradation (2) (2 blocks missing 2/4) |
| M00368 Ethylene biosynthesis, methionine => ethylene (1) (2 blocks missing 1/3) | M00368 Ethylene biosynthesis, methionine => ethylene (1) (2 blocks missing 1/3) | M00368 Ethylene biosynthesis, methionine => ethylene (1) (2 blocks missing 1/3) |
| Branched-chain amino acid metabolism | Branched-chain amino acid metabolism | Branched-chain amino acid metabolism |
| M00019 Valine/isoleucine biosynthesis, pyruvate => valine / 2-oxobutanoate => isoleucine (5) (complete 4/4) | M00019 Valine/isoleucine biosynthesis, pyruvate => valine / 2-oxobutanoate => isoleucine (5) (complete 4/4) | M00019 Valine/isoleucine biosynthesis, pyruvate => valine / 2-oxobutanoate => isoleucine (5) (complete 4/4) |
| M00535 Isoleucine biosynthesis, pyruvate => 2-oxobutanoate (3) (1 block missing 2/3) | M00535 Isoleucine biosynthesis, pyruvate => 2-oxobutanoate (3) (1 block missing 2/3) | M00535 Isoleucine biosynthesis, pyruvate => 2-oxobutanoate (2) (2 blocks missing 1/3) |
| M00570 Isoleucine biosynthesis, threonine => 2-oxobutanoate => isoleucine (6) (complete 5/5) | M00570 Isoleucine biosynthesis, threonine => 2-oxobutanoate => isoleucine (6) (complete 5/5) | M00570 Isoleucine biosynthesis, threonine => 2-oxobutanoate => isoleucine (5) (1 block missing 4/5) |
| M00432 Leucine biosynthesis, 2-oxoisovalerate => 2-oxoisocaproate (4) (complete 3/3) | M00432 Leucine biosynthesis, 2-oxoisovalerate => 2-oxoisocaproate (4) (complete 3/3) | M00432 Leucine biosynthesis, 2-oxoisovalerate => 2-oxoisocaproate (3) (1 block missing 2/3) |
| M00036 Leucine degradation, leucine => acetoacetate + acetyl-CoA (2) (incomplete 1/6) | M00036 Leucine degradation, leucine => acetoacetate + acetyl-CoA (2) (incomplete 1/6) | M00036 Leucine degradation, leucine => acetoacetate + acetyl-CoA (2) (incomplete 1/6) |
| Lysine metabolism | Lysine metabolism | Lysine metabolism |
| M00016 Lysine biosynthesis, succinyl-DAP pathway, aspartate => lysine (10) (complete 9/9) | M00016 Lysine biosynthesis, succinyl-DAP pathway, aspartate => lysine (9) (complete 9/9) | M00016 Lysine biosynthesis, succinyl-DAP pathway, aspartate => lysine (9) (complete 9/9) |
| M00525 Lysine biosynthesis, acetyl-DAP pathway, aspartate => lysine (6) (incomplete 6/9) | M00525 Lysine biosynthesis, acetyl-DAP pathway, aspartate => lysine (6) (incomplete 6/9) | M00525 Lysine biosynthesis, acetyl-DAP pathway, aspartate => lysine (6) (incomplete 6/9) |
| M00526 Lysine biosynthesis, DAP dehydrogenase pathway, aspartate => lysine (5) (1 block missing 5/6) | M00526 Lysine biosynthesis, DAP dehydrogenase pathway, aspartate => lysine (5) (1 block missing 5/6) | M00526 Lysine biosynthesis, DAP dehydrogenase pathway, aspartate => lysine (5) (1 block missing 5/6) |
| M00527 Lysine biosynthesis, DAP aminotransferase pathway, aspartate => lysine (6) (1 block missing 6/7) | M00527 Lysine biosynthesis, DAP aminotransferase pathway, aspartate => lysine (6) (1 block missing 6/7) | M00527 Lysine biosynthesis, DAP aminotransferase pathway, aspartate => lysine (6) (1 block missing 6/7) |
| M00956 Lysine degradation, bacteria, L-lysine => succinate (1) (incomplete 1/7) | ABSENT | ABSENT |
| M00957 Lysine degradation, bacteria, L-lysine => glutarate => succinate/acetyl-CoA (1) (incomplete 1/5) | ABSENT | ABSENT |

| | | |
|---|---|---|
| M00960 Lysine degradation, bacteria, L-lysine => D-lysine => succinate (2) (incomplete 2/9) | M00960 Lysine degradation, bacteria, L-lysine => D-lysine => succinate (1) (incomplete 1/9) | M00960 Lysine degradation, bacteria, L-lysine => D-lysine => succinate (1) (incomplete 1/9) |
| Arginine and proline metabolism | Arginine and proline metabolism | Arginine and proline metabolism |
| M00028 Ornithine biosynthesis, glutamate => ornithine (5) (complete 4/4) | M00028 Ornithine biosynthesis, glutamate => ornithine (4) (complete 4/4) | M00028 Ornithine biosynthesis, glutamate => ornithine (4) (complete 4/4) |
| M00844 Arginine biosynthesis, ornithine => arginine (3) (complete 3/3) | M00844 Arginine biosynthesis, ornithine => arginine (3) (complete 3/3) | M00844 Arginine biosynthesis, ornithine => arginine (3) (complete 3/3) |
| M00845 Arginine biosynthesis, glutamate => acetylcitrulline => arginine (5) (2 blocks missing 5/7) | M00845 Arginine biosynthesis, glutamate => acetylcitrulline => arginine (4) (incomplete 4/7) | M00845 Arginine biosynthesis, glutamate => acetylcitrulline => arginine (4) (incomplete 4/7) |
| M00029 Urea cycle (4) (1 block missing 4/5) | M00029 Urea cycle (3) (2 blocks missing 3/5) | M00029 Urea cycle (3) (2 blocks missing 3/5) |
| M00015 Proline biosynthesis, glutamate => proline (3) (complete 2/2) | M00015 Proline biosynthesis, glutamate => proline (3) (complete 2/2) | M00015 Proline biosynthesis, glutamate => proline (3) (complete 2/2) |
| M00970 Proline degradation, proline => glutamate (1) (complete 1/1) | M00970 Proline degradation, proline => glutamate (1) (complete 1/1) | **ABSENT** |
| M00972 Proline metabolism (1) (2 blocks missing 1/3) | M00972 Proline metabolism (1) (2 blocks missing 1/3) | M00972 Proline metabolism (1) (2 blocks missing 1/3) |
| Polyamine biosynthesis | Polyamine biosynthesis | Polyamine biosynthesis |
| M00133 Polyamine biosynthesis, arginine => agmatine => putrescine => spermidine (2) (2 blocks missing 2/4) | M00133 Polyamine biosynthesis, arginine => agmatine => putrescine => spermidine (2) (2 blocks missing 2/4) | M00133 Polyamine biosynthesis, arginine => agmatine => putrescine => spermidine (2) (2 blocks missing 2/4) |
| M00134 Polyamine biosynthesis, arginine => ornithine => putrescine (2) (complete 2/2) | M00134 Polyamine biosynthesis, arginine => ornithine => putrescine (1) (1 block missing 1/2) | M00134 Polyamine biosynthesis, arginine => ornithine => putrescine (1) (1 block missing 1/2) |
| M00136 GABA biosynthesis, prokaryotes, putrescine => GABA (1) (incomplete 1/4) | | |
| Histidine metabolism | Histidine metabolism | Histidine metabolism |
| M00026 Histidine biosynthesis, PRPP => histidine (9) (1 block missing 5/6) | M00026 Histidine biosynthesis, PRPP => histidine (11) (complete 6/6) | M00026 Histidine biosynthesis, PRPP => histidine (6) (incomplete 2/6) |
| M00045 Histidine degradation, histidine => N-formiminoglutamate => glutamate (5) (complete 4/4) | **ABSENT** | **ABSENT** |
| Aromatic amino acid metabolism | Aromatic amino acid metabolism | Aromatic amino acid metabolism |
| M00022 Shikimate pathway, phosphoenolpyruvate + erythrose-4P => chorismate (6) (complete 4/4) | M00022 Shikimate pathway, phosphoenolpyruvate + erythrose-4P => chorismate (6) (complete 4/4) | M00022 Shikimate pathway, phosphoenolpyruvate + erythrose-4P => chorismate (6) (complete 4/4) |
| M00023 Tryptophan biosynthesis, chorismate => tryptophan (7) (complete 3/3) | M00023 Tryptophan biosynthesis, chorismate => tryptophan (7) (complete 3/3) | M00023 Tryptophan biosynthesis, chorismate => tryptophan (7) (complete 3/3) |
| M00024 Phenylalanine biosynthesis, chorismate => phenylpyruvate => phenylalanine (2) (1 block missing 1/2) | M00024 Phenylalanine biosynthesis, chorismate => phenylpyruvate => phenylalanine (2) (1 block missing 1/2) | M00024 Phenylalanine biosynthesis, chorismate => phenylpyruvate => phenylalanine (2) (1 block missing 1/2) |
| M00040 Tyrosine biosynthesis, chorismate => arogenate => tyrosine (2) (1 block missing 2/3) | M00040 Tyrosine biosynthesis, chorismate => arogenate => tyrosine (2) (1 block missing 2/3) | M00040 Tyrosine biosynthesis, chorismate => arogenate => tyrosine (2) (1 block missing 2/3) |
| M00042 Catecholamine biosynthesis, tyrosine => dopamine => noradrenaline => adrenaline (1) (incomplete 1/4) | **ABSENT** | **ABSENT** |
| M00037 Melatonin biosynthesis, animals, tryptophan => serotonin => melatonin (1) (incomplete 1/4) | M00037 Melatonin biosynthesis, animals, tryptophan => serotonin => melatonin (1) (incomplete 1/4) | **ABSENT** |
| M00936 Melatonin biosynthesis, plants, tryptophan => serotonin => melatonin (1) (incomplete 1/4) | **ABSENT** | **ABSENT** |
| Other amino acid metabolism | Other amino acid metabolism | Other amino acid metabolism |
| M00027 GABA (gamma-Aminobutyrate) shunt (1) (2 blocks missing 1/3) | **ABSENT** | **ABSENT** |
| M00118 Glutathione biosynthesis, glutamate => glutathione (2) (complete 2/2) | M00118 Glutathione biosynthesis, glutamate => glutathione (2) (complete 2/2) | M00118 Glutathione biosynthesis, glutamate => glutathione (2) (complete 2/2) |
| **Glycan metabolism** | **Glycan metabolism** | **Glycan metabolism** |

| | | |
|---|---|---|
| Glycosaminoglycan metabolism | ABSENT | ABSENT |
| M00079 Keratan sulfate degradation (1) (incomplete 1/4) | | |
| Lipopolysaccharide metabolism | Lipopolysaccharide metabolism | Lipopolysaccharide metabolism |
| M00060 KDO2-lipid A biosynthesis, Raetz pathway, LpxL-LpxM type (8) (1 block missing 8/9) | M00060 KDO2-lipid A biosynthesis, Raetz pathway, LpxL-LpxM type (8) (1 block missing 8/9) | M00060 KDO2-lipid A biosynthesis, Raetz pathway, LpxL-LpxM type (8) (1 block missing 8/9) |
| M00866 KDO2-lipid A biosynthesis, Raetz pathway, non-LpxL-LpxM type (8) (1 block missing 8/9) | M00866 KDO2-lipid A biosynthesis, Raetz pathway, non-LpxL-LpxM type (8) (1 block missing 8/9) | M00866 KDO2-lipid A biosynthesis, Raetz pathway, non-LpxL-LpxM type (8) (1 block missing 8/9) |
| M00867 KDO2-lipid A modification pathway (1) (incomplete 1/5) | M00867 KDO2-lipid A modification pathway (1) (incomplete 1/5) | M00867 KDO2-lipid A modification pathway (1) (incomplete 1/5) |
| M00063 CMP-KDO biosynthesis (3) (1 block missing 3/4) | M00063 CMP-KDO biosynthesis (3) (1 block missing 3/4) | M00063 CMP-KDO biosynthesis (3) (1 block missing 3/4) |
| M00064 ADP-L-glycero-D-manno-heptose biosynthesis (2) (2 blocks missing 3/5) | M00064 ADP-L-glycero-D-manno-heptose biosynthesis (3) (1 block missing 4/5) | M00064 ADP-L-glycero-D-manno-heptose biosynthesis (3) (1 block missing 4/5) |
| **Metabolism of cofactors and vitamins** | **Metabolism of cofactors and vitamins** | **Metabolism of cofactors and vitamins** |
| Cofactor and vitamin metabolism | Cofactor and vitamin metabolism | Cofactor and vitamin metabolism |
| M00127 Thiamine biosynthesis, prokaryotes, AIR (+ DXP/tyrosine) => TMP/TPP (6) (incomplete 4/7) | M00127 Thiamine biosynthesis, prokaryotes, AIR (+ DXP/tyrosine) => TMP/TPP (6) (incomplete 4/7) | M00127 Thiamine biosynthesis, prokaryotes, AIR (+ DXP/tyrosine) => TMP/TPP (6) (incomplete 4/7) |
| M00895 Thiamine biosynthesis, prokaryotes, AIR (+ DXP/glycine) => TMP/TPP (7) (incomplete 6/9) | M00895 Thiamine biosynthesis, prokaryotes, AIR (+ DXP/glycine) => TMP/TPP (7) (incomplete 6/9) | M00895 Thiamine biosynthesis, prokaryotes, AIR (+ DXP/glycine) => TMP/TPP (7) (incomplete 6/9) |
| M00896 Thiamine biosynthesis, archaea, AIR (+ NAD+) => TMP/TPP (4) (1 block missing 3/4) | M00896 Thiamine biosynthesis, archaea, AIR (+ NAD+) => TMP/TPP (4) (1 block missing 3/4) | M00896 Thiamine biosynthesis, archaea, AIR (+ NAD+) => TMP/TPP (4) (1 block missing 3/4) |
| M00897 Thiamine biosynthesis, plants, AIR (+ NAD+) => TMP/thiamine/TPP (1) (incomplete 1/5) | M00897 Thiamine biosynthesis, plants, AIR (+ NAD+) => TMP/thiamine/TPP (1) (incomplete 1/5) | M00897 Thiamine biosynthesis, plants, AIR (+ NAD+) => TMP/thiamine/TPP (1) (incomplete 1/5) |
| M00899 Thiamine salvage pathway, HMP/HET => TMP (2) (1 block missing 1/2) | M00899 Thiamine salvage pathway, HMP/HET => TMP (2) (1 block missing 1/2) | M00899 Thiamine salvage pathway, HMP/HET => TMP (2) (1 block missing 1/2) |
| M00125 Riboflavin biosynthesis, plants and bacteria, GTP => riboflavin/FMN/FAD (5) (1 block missing 6/7) | M00125 Riboflavin biosynthesis, plants and bacteria, GTP => riboflavin/FMN/FAD (5) (1 block missing 6/7) | M00125 Riboflavin biosynthesis, plants and bacteria, GTP => riboflavin/FMN/FAD (5) (1 block missing 6/7) |
| M00911 Riboflavin biosynthesis, fungi, GTP => riboflavin/FMN/FAD (2) (incomplete 2/8) | M00911 Riboflavin biosynthesis, fungi, GTP => riboflavin/FMN/FAD (2) (incomplete 2/8) | M00911 Riboflavin biosynthesis, fungi, GTP => riboflavin/FMN/FAD (2) (incomplete 2/8) |
| M00124 Pyridoxal-P biosynthesis, erythrose-4P => pyridoxal-P (4) (2 blocks missing 4/6) | M00124 Pyridoxal-P biosynthesis, erythrose-4P => pyridoxal-P (4) (2 blocks missing 4/6) | M00124 Pyridoxal-P biosynthesis, erythrose-4P => pyridoxal-P (3) (incomplete 3/6) |
| M00115 NAD biosynthesis, aspartate => quinolinate => NAD (5) (complete 5/5) | M00115 NAD biosynthesis, aspartate => quinolinate => NAD (4) (1 block missing 4/5) | M00115 NAD biosynthesis, aspartate => quinolinate => NAD (5) (complete 5/5) |
| M00912 NAD biosynthesis, tryptophan => quinolinate => NAD (3) (incomplete 3/8) | M00912 NAD biosynthesis, tryptophan => quinolinate => NAD (2) (incomplete 2/8) | M00912 NAD biosynthesis, tryptophan => quinolinate => NAD (3) (incomplete 3/8) |
| M00119 Pantothenate biosynthesis, valine/L-aspartate => pantothenate (3) (2 blocks missing 3/5) | M00119 Pantothenate biosynthesis, valine/L-aspartate => pantothenate (1) (incomplete 1/5) | M00119 Pantothenate biosynthesis, valine/L-aspartate => pantothenate (1) (incomplete 1/5) |
| M00913 Pantothenate biosynthesis, 2-oxoisovalerate/spermine => pantothenate (2) (incomplete 2/5) | ABSENT | ABSENT |
| M00120 Coenzyme A biosynthesis, pantothenate => CoA (4) (complete 3/3) | M00120 Coenzyme A biosynthesis, pantothenate => CoA (4) (complete 3/3) | M00120 Coenzyme A biosynthesis, pantothenate => CoA (4) (complete 3/3) |
| M00914 Coenzyme A biosynthesis, archaea, 2-oxoisovalerate => 4-phosphopantoate => CoA (2) (incomplete 2/7) | M00914 Coenzyme A biosynthesis, archaea, 2-oxoisovalerate => 4-phosphopantoate => CoA (1) (incomplete 1/7) | M00914 Coenzyme A biosynthesis, archaea, 2-oxoisovalerate => 4-phosphopantoate => CoA (1) (incomplete 1/7) |
| M00572 Pimeloyl-ACP biosynthesis, BioC-BioH pathway, malonyl-ACP => pimeloyl-ACP (6) (complete 6/6) | M00572 Pimeloyl-ACP biosynthesis, BioC-BioH pathway, malonyl-ACP => pimeloyl-ACP (6) (complete 6/6) | M00572 Pimeloyl-ACP biosynthesis, BioC-BioH pathway, malonyl-ACP => pimeloyl-ACP (5) (1 block missing 5/6) |
| M00123 Biotin biosynthesis, pimeloyl-ACP/CoA => biotin (4) (complete 3/3) | M00123 Biotin biosynthesis, pimeloyl-ACP/CoA => biotin (4) (complete 3/3) | M00123 Biotin biosynthesis, pimeloyl-ACP/CoA => biotin (3) (1 block missing 2/3) |
| M00950 Biotin biosynthesis, BioU pathway, pimeloyl-ACP/CoA => biotin (3) (1 block missing 3/4) | M00950 Biotin biosynthesis, BioU pathway, pimeloyl-ACP/CoA => biotin (3) (1 block missing 3/4) | M00950 Biotin biosynthesis, BioU pathway, pimeloyl-ACP/CoA => biotin (2) (2 blocks missing 2/4) |

| | | |
|---|---|---|
| M00573 Biotin biosynthesis, BioI pathway, long-chain-acyl-ACP => pimeloyl-ACP => biotin (3) (2 blocks missing 3/5) | M00573 Biotin biosynthesis, BioI pathway, long-chain-acyl-ACP => pimeloyl-ACP => biotin (3) (2 blocks missing 3/5) | M00573 Biotin biosynthesis, BioI pathway, long-chain-acyl-ACP => pimeloyl-ACP => biotin (2) (incomplete 2/5) |
| M00577 Biotin biosynthesis, BioW pathway, pimelate => pimeloyl-CoA => biotin (4) (1 block missing 4/5) | M00577 Biotin biosynthesis, BioW pathway, pimelate => pimeloyl-CoA => biotin (4) (1 block missing 4/5) | M00577 Biotin biosynthesis, BioW pathway, pimelate => pimeloyl-CoA => biotin (3) (2 blocks missing 3/5) |
| M00881 Lipoic acid biosynthesis, plants and bacteria, octanoyl-ACP => dihydrolipoyl-E2/H (2) (complete 2/2) | M00881 Lipoic acid biosynthesis, plants and bacteria, octanoyl-ACP => dihydrolipoyl-E2/H (2) (complete 2/2) | M00881 Lipoic acid biosynthesis, plants and bacteria, octanoyl-ACP => dihydrolipoyl-E2/H (2) (complete 2/2) |
| M00882 Lipoic acid biosynthesis, eukaryotes, octanoyl-ACP => dihydrolipoyl-H (1) (1 block missing 1/2) | M00882 Lipoic acid biosynthesis, eukaryotes, octanoyl-ACP => dihydrolipoyl-H (1) (1 block missing 1/2) | M00882 Lipoic acid biosynthesis, eukaryotes, octanoyl-ACP => dihydrolipoyl-H (1) (1 block missing 1/2) |
| M00883 Lipoic acid biosynthesis, animals and bacteria, octanoyl-ACP => dihydrolipoyl-H => dihydrolipoyl-E2 (1) (2 blocks missing 1/3) | M00883 Lipoic acid biosynthesis, animals and bacteria, octanoyl-ACP => dihydrolipoyl-H => dihydrolipoyl-E2 (1) (2 blocks missing 1/3) | M00883 Lipoic acid biosynthesis, animals and bacteria, octanoyl-ACP => dihydrolipoyl-H => dihydrolipoyl-E2 (1) (2 blocks missing 1/3) |
| M00884 Lipoic acid biosynthesis, octanoyl-CoA => dihydrolipoyl-E2 (1) (1 block missing 1/2) | M00884 Lipoic acid biosynthesis, octanoyl-CoA => dihydrolipoyl-E2 (1) (1 block missing 1/2) | M00884 Lipoic acid biosynthesis, octanoyl-CoA => dihydrolipoyl-E2 (1) (1 block missing 1/2) |
| M00126 Tetrahydrofolate biosynthesis, GTP => THF (4) (2 blocks missing 3/5) | M00126 Tetrahydrofolate biosynthesis, GTP => THF (4) (2 blocks missing 3/5) | M00126 Tetrahydrofolate biosynthesis, GTP => THF (4) (2 blocks missing 3/5) |
| M00840 Tetrahydrofolate biosynthesis, mediated by ribA and trpF, GTP => THF (1) (incomplete 1/6) | M00840 Tetrahydrofolate biosynthesis, mediated by ribA and trpF, GTP => THF (1) (incomplete 1/6) | M00840 Tetrahydrofolate biosynthesis, mediated by ribA and trpF, GTP => THF (1) (incomplete 1/6) |
| M00841 Tetrahydrofolate biosynthesis, mediated by PTPS, GTP => THF (2) (incomplete 2/5) | M00841 Tetrahydrofolate biosynthesis, mediated by PTPS, GTP => THF (2) (incomplete 2/5) | M00841 Tetrahydrofolate biosynthesis, mediated by PTPS, GTP => THF (2) (incomplete 2/5) |
| M00842 Tetrahydrobiopterin biosynthesis, GTP => BH4 (2) (1 block missing 2/3) | M00842 Tetrahydrobiopterin biosynthesis, GTP => BH4 (2) (1 block missing 2/3) | M00842 Tetrahydrobiopterin biosynthesis, GTP => BH4 (2) (1 block missing 2/3) |
| M00843 L-threo-Tetrahydrobiopterin biosynthesis, GTP => L-threo-BH4 (2) (1 block missing 2/3) | M00843 L-threo-Tetrahydrobiopterin biosynthesis, GTP => L-threo-BH4 (2) (1 block missing 2/3) | M00843 L-threo-Tetrahydrobiopterin biosynthesis, GTP => L-threo-BH4 (2) (1 block missing 2/3) |
| M00880 Molybdenum cofactor biosynthesis, GTP => molybdenum cofactor (4) (1 block missing 2/3) | **ABSENT** | **ABSENT** |
| M00140 C1-unit interconversion, prokaryotes (2) (1 block missing 2/3) | M00140 C1-unit interconversion, prokaryotes (2) (1 block missing 2/3) | M00140 C1-unit interconversion, prokaryotes (2) (1 block missing 2/3) |
| M00141 C1-unit interconversion, eukaryotes (1) (1 block missing 1/2) | M00141 C1-unit interconversion, eukaryotes (1) (1 block missing 1/2) | M00141 C1-unit interconversion, eukaryotes (1) (1 block missing 1/2) |
| M00846 Siroheme biosynthesis, glutamyl-tRNA => siroheme (4) (2 blocks missing 4/6) | M00846 Siroheme biosynthesis, glutamyl-tRNA => siroheme (4) (2 blocks missing 4/6) | M00846 Siroheme biosynthesis, glutamyl-tRNA => siroheme (4) (2 blocks missing 4/6) |
| M00868 Heme biosynthesis, animals and fungi, glycine => heme (7) (1 block missing 7/8) | M00868 Heme biosynthesis, animals and fungi, glycine => heme (7) (1 block missing 7/8) | M00868 Heme biosynthesis, animals and fungi, glycine => heme (7) (1 block missing 7/8) |
| M00121 Heme biosynthesis, plants and bacteria, glutamate => heme (8) (2 blocks missing 8/10) | M00121 Heme biosynthesis, plants and bacteria, glutamate => heme (8) (2 blocks missing 8/10) | M00121 Heme biosynthesis, plants and bacteria, glutamate => heme (8) (2 blocks missing 8/10) |
| M00926 Heme biosynthesis, bacteria, glutamyl-tRNA => coproporphyrin III => heme (5) (incomplete 5/9) | M00926 Heme biosynthesis, bacteria, glutamyl-tRNA => coproporphyrin III => heme (5) (incomplete 5/9) | M00926 Heme biosynthesis, bacteria, glutamyl-tRNA => coproporphyrin III => heme (5) (incomplete 5/9) |
| M00924 Cobalamin biosynthesis, anaerobic, uroporphyrinogen III => sirohydrochlorin => cobyrinate a,c-diamide (9) (2 blocks missing 9/11) | M00924 Cobalamin biosynthesis, anaerobic, uroporphyrinogen III => sirohydrochlorin => cobyrinate a,c-diamide (9) (2 blocks missing 9/11) | M00924 Cobalamin biosynthesis, anaerobic, uroporphyrinogen III => sirohydrochlorin => cobyrinate a,c-diamide (6) (incomplete 6/11) |
| M00925 Cobalamin biosynthesis, aerobic, uroporphyrinogen III => precorrin 2 => cobyrinate a,c-diamide (11) (2 blocks missing 9/11) | M00925 Cobalamin biosynthesis, aerobic, uroporphyrinogen III => precorrin 2 => cobyrinate a,c-diamide (11) (2 blocks missing 9/11) | M00925 Cobalamin biosynthesis, aerobic, uroporphyrinogen III => precorrin 2 => cobyrinate a,c-diamide (8) (incomplete 6/11) |
| M00122 Cobalamin biosynthesis, cobyrinate a,c-diamide => cobalamin (7) (2 blocks missing 5/7) | M00122 Cobalamin biosynthesis, cobyrinate a,c-diamide => cobalamin (8) (complete 7/7) | M00122 Cobalamin biosynthesis, cobyrinate a,c-diamide => cobalamin (3) (incomplete 3/7) |
| M00117 Ubiquinone biosynthesis, prokaryotes, chorismate (+ polyprenyl-PP) => ubiquinol (4) (incomplete 5/9) | M00117 Ubiquinone biosynthesis, prokaryotes, chorismate (+ polyprenyl-PP) => ubiquinol (4) (incomplete 5/9) | M00117 Ubiquinone biosynthesis, prokaryotes, chorismate (+ polyprenyl-PP) => ubiquinol (4) (incomplete 5/9) |
| M00116 Menaquinone biosynthesis, chorismate (+ polyprenyl-PP) => menaquinol (1) (incomplete 1/9) | M00116 Menaquinone biosynthesis, chorismate (+ polyprenyl-PP) => menaquinol (1) (incomplete 1/9) | M00116 Menaquinone biosynthesis, chorismate (+ polyprenyl-PP) => menaquinol (1) (incomplete 1/9) |
| **Biosynthesis of terpenoids and polyketides** | **Biosynthesis of terpenoids and polyketides** | **Biosynthesis of terpenoids and polyketides** |
| Terpenoid backbone biosynthesis | Terpenoid backbone biosynthesis | Terpenoid backbone biosynthesis |
| M00096 C5 isoprenoid biosynthesis, non-mevalonate pathway (6) (1 block missing 7/8) | M00096 C5 isoprenoid biosynthesis, non-mevalonate pathway (6) (1 block missing 7/8) | M00096 C5 isoprenoid biosynthesis, non-mevalonate pathway (6) (1 block missing 7/8) |

| | | |
|---|---|---|
| M00364 C10-C20 isoprenoid biosynthesis, bacteria (1) (1 block missing 1/2) | M00364 C10-C20 isoprenoid biosynthesis, bacteria (1) (1 block missing 1/2) | M00364 C10-C20 isoprenoid biosynthesis, bacteria (1) (1 block missing 1/2) |
| Polyketide sugar unit biosynthesis | Polyketide sugar unit biosynthesis | Polyketide sugar unit biosynthesis |
| M00793 dTDP-L-rhamnose biosynthesis (4) (complete 3/3) | M00793 dTDP-L-rhamnose biosynthesis (4) (complete 3/3) | M00793 dTDP-L-rhamnose biosynthesis (4) (complete 3/3) |
| **Biosynthesis of other secondary metabolites** | **Biosynthesis of other secondary metabolites** | **Biosynthesis of other secondary metabolites** |
| Biosynthesis of phytochemical compounds | Biosynthesis of phytochemical compounds | Biosynthesis of phytochemical compounds |
| M00953 Mugineic acid biosynthesis, methionine => 3-epihydroxymugineic acid (1) (incomplete 1/5) | M00953 Mugineic acid biosynthesis, methionine => 3-epihydroxymugineic acid (1) (incomplete 1/5) | M00953 Mugineic acid biosynthesis, methionine => 3-epihydroxymugineic acid (1) (incomplete 1/5) |
| **Xenobiotics biodegradation** | **Xenobiotics biodegradation** | **Xenobiotics biodegradation** |
| Aromatics degradation | Aromatics degradation | Aromatics degradation |
| M00569 Catechol meta-cleavage, catechol => acetyl-CoA / 4-methylcatechol => propanoyl-CoA (1) (incomplete 1/5) | **ABSENT** | **ABSENT** |
| M00878 Phenylacetate degradation, phenylaxetate => acetyl-CoA/succinyl-CoA (1) (incomplete 1/7) | M00878 Phenylacetate degradation, phenylaxetate => acetyl-CoA/succinyl-CoA (1) (incomplete 1/7) | M00878 Phenylacetate degradation, phenylaxetate => acetyl-CoA/succinyl-CoA (1) (incomplete 1/7) |
| M00545 Trans-cinnamate degradation, trans-cinnamate => acetyl-CoA (1) (incomplete 1/6) | **ABSENT** | **ABSENT** |
| **Gene set** | **Gene set** | **Gene set** |
| Pathogenicity | Pathogenicity | Pathogenicity |
| M00575 Pertussis pathogenicity signature, T1SS (3) (2 blocks missing 3/5) | M00575 Pertussis pathogenicity signature, T1SS (3) (2 blocks missing 3/5) | M00575 Pertussis pathogenicity signature, T1SS (1) (incomplete 1/5) |
| Drug resistance | Drug resistance | Drug resistance |
| M00627 beta-Lactam resistance, Bla system (1) (2 blocks missing 1/3) | M00627 beta-Lactam resistance, Bla system (1) (2 blocks missing 1/3) | **ABSENT** |
| M00726 Cationic antimicrobial peptide (CAMP) resistance, lysyl-phosphatidylglycerol (L-PG) synthase MprF (1) (2 blocks missing 1/3) | M00726 Cationic antimicrobial peptide (CAMP) resistance, lysyl-phosphatidylglycerol (L-PG) synthase MprF (1) (2 blocks missing 1/3) | M00726 Cationic antimicrobial peptide (CAMP) resistance, lysyl-phosphatidylglycerol (L-PG) synthase MprF (1) (2 blocks missing 1/3) |
| M00718 Multidrug resistance, efflux pump MexAB-OprM (3) (1 block missing 1/2) | M00718 Multidrug resistance, efflux pump MexAB-OprM (3) (1 block missing 1/2) | **ABSENT** |
| M00642 Multidrug resistance, efflux pump MexJK-OprM (2) (1 block missing 1/2) | **ABSENT** | **ABSENT** |
| **Module set** | | |
| Metabolic capacity | **ABSENT** | **ABSENT** |
| M00618 Acetogen (0) (1 block missing 1/2) | | |
| **Legend:** | | |
| pathway present in Asaia bogorensis genom and absent in *Ca.* Kirkpatrella diaphorinas and *Acetobacteraceae*_TRYOCC genomes | | |
| pathway present in Asaia bogorensis and *Ca.* Kirkpatrella diaphorinas genomes but absent in *Acetobacteraceae*_TRYOCC genome | | |
| pathway with missing gne(s) in *Acetobacteraceae*_TRYOCC relative to *Ca.* Kirkpatrella diaphorinas | | |

**Table S1.** The comparison of metabolic pathways of *Asia bogorensis W19*, *Kirkpatrickella* and *Acetobacteraceae TRYOCC.*

| **Fluorescence *in situ* hybridization (FISH)** | | | | | |
|---|---|---|---|---|---|
| **Probe name** | **Probe sequence** | **Specificity** | **Fluorophore** | **Hybridization buffer** | **References** |
| Sul664R | CCMCACATTCCAGYTACTCC | 16S rRNA gene of *Sulcia* | Cy3 | 1M Tris-HCl (pH 8.0) – 1 ml, 5M NaCl – 9 ml, 20% SDS – 20 µl, 30% formamide – 15 ml, water – up to 50 ml | Koga et al., 2013 |
| Bet940R | TTAATCCACATCATCCACCG | 16S rRNA gene of *Vidania* | AlexaFluor 488 | | Demanèche et al. 2008 |
| *Asaia* 1 | ATAACATCGGGAAACTGGTGCT | 16S rRNA gene of *Acetobacteraceae* symbiont | Cy5 | | Favia et al., 2007 |
| *Asaia* 1 | GAAATACCCATCTCTGGATA | 16S rRNA gene of *Acetobacteraceae* symbiont | Cy5 | | Favia et al., 2007 |

**Table S2.** Sequences of symbiont-specific fluorochrome-labelled oligonucleotide probes.

| **Genome** | **Bioproject** | **Biosample** | **Assembly** |
|---|---|---|---|
| *Acetobacter pomorum* | PRJNA397225 | SAMN07682859 | GCA_002456135.1 |
| *Acetobacter tropicalis DM* | PRJNA397225 | SAMN07452474 | GCA_002549835.1 |
| *Acetobacter tropicalis* | PRJNA642854 | SAMN15399467 | GCA_014486685.1 |
| *Acetobacter aceti* | PRJNA311264 | SAMN04396917 | GCA_002005445.1 |
| *Gluconacetobacter johannae* | PRJNA627338 | SAMN14650889 | GCA_014174305.1 |
| *Gluconacetobacter liquefaciens* | PRJNA627338 | SAMN14766534 | GCA_014174285.1 |
| *Gluconacetobacter diazotrophicus* | PRJNA21071 | SAMN02598444 | GCA_000021325.1 |
| *Asaia platycodi* | PRJDB613 | SAMD00002827 | GCA_000614545.1 |
| *Asaia prunellae* | PRJDB589 | SAMD00004065 | GCA_000613885.1 |
| *Asaia siamensis* | PRJDB10511 | SAMD00244847 | GCA_014635085.1 |
| *Asaia astilbis* | PRJDB587 | SAMD00009367 | GCA_000613845.1 |
| *Asaia bogorensis OTPB* | PRJDB519 | SAMD00019081 | GCA_001547995.1 |
| *Asaia* sp. SF2 | PRJNA219210 | SAMN02469759 | GCA_000505765.1 |
| *Asaia lannensis* | PRJNA849918 | SAMN29145435 | GCA_024054035.1 |
| *Asaia bogorensis MRS* | PRJNA756835 | SAMN20927460 | GCA_019823045.1 |
| *Asaia bogorensis* W19 | PRJNA427835 | SAMN08274829 | GCA_003994335.1 |
| *Ca.* Kirkpatrickella diaphorinae | PRJNA880923 | SAMN30877677 | GCA_025736875.1 |
| *Kozakia baliensis* | PRJNA311264 | SAMN04396915 | GCF_001787355.1 |
| *Neoasaia chiangmaiensis* | PRJNA311264 | SAMN04396918 | GCA_002005465.1 |
| *Acetobacter methanolica* | PRJNA500315 | SAMN10363313 | GCA_004346035.1 |
| *Saccharibacter floricola* | PRJNA181373 | SAMN02440467 | GCA_000378165.1 |
| *Saccharibacter* sp. 17.LH.SD | PRJNA596238 | SAMN13615857 | GCA_009834805.1 |
| *Acetobacteraceae* bacterium strain 880 chromosome | PRJNA532976 | SAMN11430874 | CP039459.1 |
| *Acetobacteraceae* bacterium strain 868 chromosome | PRJNA532976 | SAMN11430873 | CP039460.1 |
| *Gluconobacter oxydans* | PRJNA188081 | SAMN05771126 | GCA_000583855.1 |
| *Gluconobacter morbifer* | PRJNA73361 | SAMN02470912 | GCA_000234355.2 |

**Table S3.** Accession numbers of *Acetobacteraceae* genomes used for phylogenomic analysis.
