## Supplementary figures and images for "Facultatively intra-bacterial localization of a planthopper endosymbiont as an adaptation to its vertical transmission"

### Supplemental figure S1

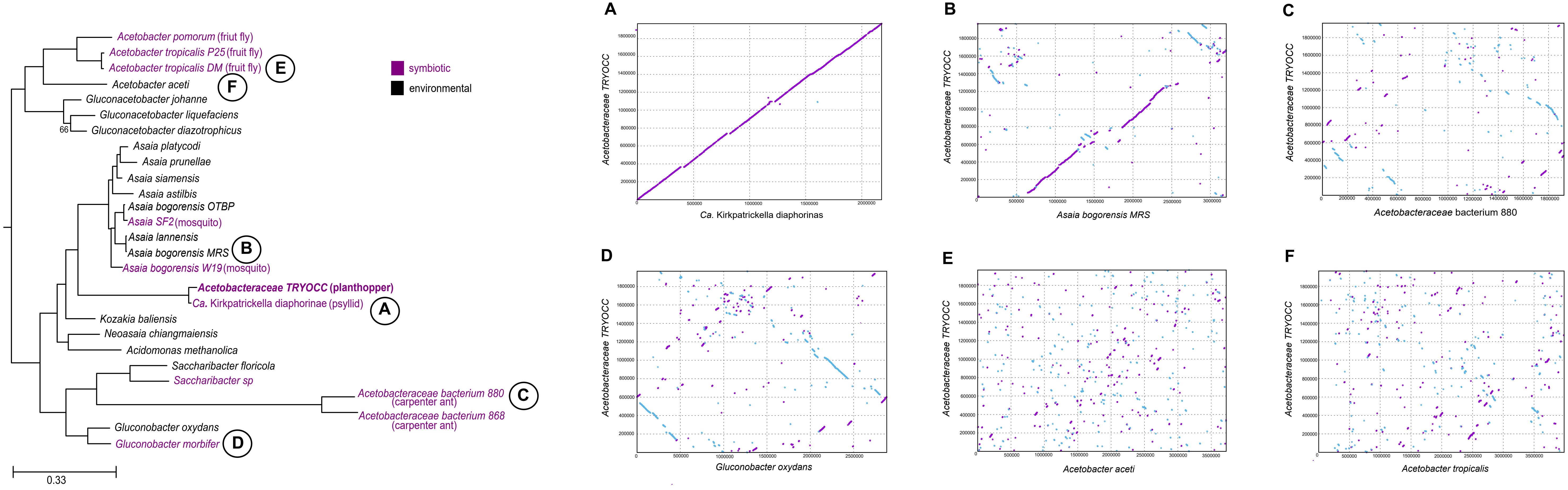
